# A humanized neuronal model system reveals key roles for manganese in neuronal endocytosis, calcium flux and mitochondrial bioenergetics

**DOI:** 10.1101/2024.11.27.625600

**Authors:** Dimitri Budinger, Sharmin Alhaque, Ramón González-Méndez, Chris Dadswell, Katy Barwick, Arianna Ferrini, Charlotte Roth, Conor J. McCann, Karin Tuschl, Fatma Al Jasmi, Maha S. Zaki, Julien H. Park, Russell C. Dale, Shekeeb Mohammad, John Christodoulou, Dale Moulding, Michael R. Duchen, Serena Barral, Manju A. Kurian

## Abstract

Manganese (Mn) is an essential trace metal that is necessary for life. Its duality as both a crucial micronutrient and potential neurotoxicant necessitates tight control of intracellular and extracellular Mn levels. Dysregulation of Mn is implicated in a broad range of human diseases, from neurodevelopmental sequelae related to Mn levels in drinking water, to acquired forms of manganism, rare inherited Mn transportopathies and more common disorders such as Parkinson’s and Alzheimer’s disease. Despite the clear association between Mn dysregulation and neurodevelopmental or neurodegenerative diseases, the underlying cellular mechanisms that govern neuropathology remain poorly understood. We established an induced pluripotent stem cells-derived midbrain neuronal system from SLC39A14, SLC39A8, and SLC30A10 patients to better understand the neuronal sequelae of Mn dysregulation. By integrating transcriptomic and functional approaches, we show that Mn dyshomeostasis leads to dysregulation of key cellular pathways that are crucial to normal neuronal function, including defects in mitochondrial bioenergetics, calcium signalling, endocytosis, and glycosylation, as well as cellular stress and early neurodegeneration. Our humanized model has enhanced understanding of the role of Mn in the human brain, and the consequences of both acquired and genetic disorders associated with Mn dysregulation. Better understanding of these underlying pathophysiological processes will identify potential targets for future therapeutic intervention.

## Introduction

Manganese (Mn) is an essential trace metal that is vital to all living organisms for the catalytic function of multiple metalloenzymes and many other metabolic processes. In humans, dietary intake of Mn is tightly regulated through the enterohepatic circulation, hepatic metabolism and biliary excretion.^1^ Circulating red blood cells mediate systemic delivery of Mn through metal transporters that regulate the transfer of Mn across cellular membranes into target organs with high energy demand, particularly the liver, skeletal system, pancreas, kidneys, and brain.^2^ In the central nervous system (CNS), Mn additionally enters the brain via the choroid plexus, olfactory epithelium and olfactory nerves, as well as through direct intra-axonal uptake via presynaptic nerve endings.^3^ Within the CNS, homoeostatic balance is tightly controlled by transmembrane transporters (including SPCA1, SPCA2, citrate transporters, ZIP8 and ZIP14, DMT1, transferrin, ferroportin, ZnT10, DAT, and ATP13A2), which mediate cellular influx and efflux of Mn.^4,5^

Dysregulation of Mn in the brain is associated with a broad range of neurological phenotypes, implicating a fundamental role for Mn in normal brain development and neuroprotection. Elevated levels of Mn in drinking water have been linked to neurodevelopmental sequelae in children with postulated effects on behaviour, cognition, and motor function.^6–9^ Mn toxicity or ‘manganism’ from occupational exposure, ephedrone drug abuse and other iatrogenic causes lead to severe extrapyramidal phenotypes as well as cognitive and neuropsychiatric manifestations.^10–14^ Mn dyshomeostasis is also implicated in more complex neurodegenerative disorders such as Parkinson’s disease (PD)^15^, Alzheimer’s disease (AD)^16^, Huntington’s disease (HD)^17^, and amyotrophic lateral sclerosis (ALS).^18^

The fundamental role of Mn in the brain is evident in the monogenic Mn transportopathies. Recessive mutations in *SLC30A10* and *SLC39A14*, encoding the Mn transporters ZnT10 and ZIP14 respectively, lead to hypermanganesaemia with complex dystonia-parkinsonism (HMNDYT1/2, OMIM #613280 & #617013).^19–22^ Biallelic mutations in *SLC39A8*, encoding the metal transporter ZIP8, cause hypomanganesaemia and neurodevelopmental defects associated with a congenital disorder of glycosylation (SLC39A8-CDG, OMIM #616721). Affected patients manifest in infancy with reduced blood Mn levels and a severe clinical phenotype characterised by dystonia, cranial asymmetry, dwarfism, epileptic encephalopathy, and developmental delay.^23–26^ The transferrin glycosylation defect is attributed to impaired Mn-dependent β-1,4-galactosyltransferase.^27,28^

Despite its broad impact on health, effective treatments for Mn-related disorders are currently very limited.^29^ The lack of precision therapies that can either modify or cure such diseases is partly attributable to our incomplete understanding of both the intricate mechanisms that regulate Mn homeostasis and the molecular and cellular sequelae of Mn imbalance.

To address this knowledge gap, we have established an induced pluripotent stem cell (iPSC)- derived midbrain dopaminergic (mDA) neuronal model to better understand the mechanisms that drive Mn homeostasis and investigate the downstream effects of Mn imbalance in a humanised neuronal system. Using iPSC derived from patients with HMNDYT1, HMNDYT2, SLC39A8-CDG, with age-matched controls and CRISPR-corrected isogenic lines, we have studied mutant ZnT10, ZIP14 and ZIP8 Mn transporters. Our study has revealed clear disease-related phenotypes related to impaired calcium signalling, defective mitochondrial function and impaired endocytosis. Greater insight into these processes will aid the development of future precision therapies for not only these rare genetic diseases, but also for other human diseases associated with Mn dyshomeostasis.

## Results

### Generation of an iPSC-derived mDA neuronal model of ZnT10, ZIP14 and ZIP8 deficiency confirms abnormal transport of Mn

We generated iPSCs from patients with clinically and genetically confirmed HMNDYT1 (Patient 1: c.[314_322del], p.[Ala105_Pro107del]; Patient 2: c.[77T>C], p.[Leu26Pro]),^19,22^ HMNDYT2 (Patient 1: c.[1407C>G], p.[Asn469Lys]; Patient 2: c.[781-9C>G], p.[His251Profs26]),^21,39^ and SLC39A8-CDG (Patient 1: c.[338G>C], p.[Cys113Ser]; Patient 2: c.[112G>C], p.[Gly38Ala] and c.[1019T>A], p.[Ile340Asn])^23,26^ (**Figure 1A, Table S1**) using the Sendai virus reprogramming strategy.^40^ Two clones were characterised for each patient-derived line. Three isogenic control lines were generated with CRISPR-Cas9 technology^41^, using homology-directed repair (HDR) to correct single point-mutations in ZIP14 (Patient 1), ZIP8 (Patient 1) and ZnT10 (Patient 2) (**Figure S1D**). Two previously characterised age-matched control iPSC lines were utilised for this study.^35,36,38^ All iPSC lines retained structural genomic integrity as indicated by single nucleotide polymorphism (SNP) analysis and were confirmed to be pluripotent (**Figure S1A–C, Figure S1E and S1F**).

**Figure 1.**
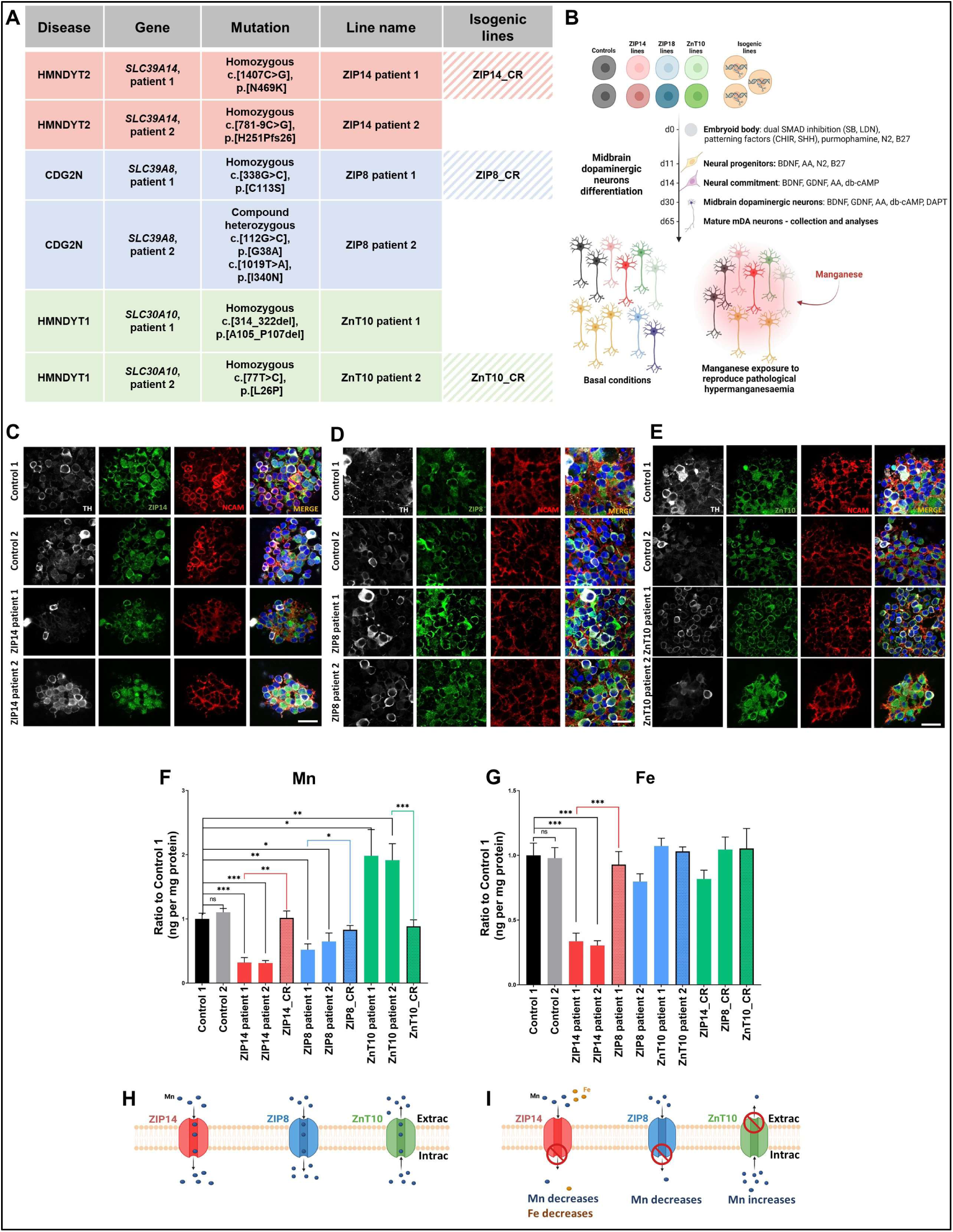
An iPSC-derived mDA neuronal system reveals normal Mn transporter expression with disease-specific impairment of Mn transport. **(A)** iPSC lines generated, corresponding genotype and line name. A total of 6 patient-derived lines were generated, together with 3 isogenic CRISPR-corrected controls. In addition, 2 previously generated iPSC age-matched control lines were also available for this study. See also **Table S1** patient information. **(B)** Workflow of iPSC generation into mDA neurons and experiments. Figure generated using BioRender. **(C-E)** Day 65 mDA neuron cultures stained for the neuron surface marker NCAM, Tyrosine hydroxylase (TH) and ZIP14 (A), ZIP8 (B), and ZnT10 (C) show expression of manganese transporters at the cell surface in control and patient lines. Scale bar, 20 µm. **(F-G)** ICP-MS analysis for intracellular levels of manganese (F) and iron (G) show manganese and iron dyshomeostasis in patient lines. N = 3 – 14 from minimum 3 independent experiments, ^∗^p=0.05-0.01, **p=0.01-0.001, p***< 0.001, unpaired Student’s t test). Values are given as means ± SEM. **(H-I)** Graphical representation of the physiological (H) and pathological (I) function of ZIP14, ZIP8, and ZnT10 in mDA neurons. Extrac; extracellular, Intrac; intracellular. Figure generated using BioRender.

Given that the midbrain, and in particular midbrain dopaminergic (mDA) neurons are exquisitely sensitive to Mn imbalance,^42–44^ we developed an iPSC-derived mDA neuronal system to study the effect of Mn dyshomeostasis in the brain (**Figure 1B**). All iPSC lines were successfully differentiated into mDA neuronal precursors with high efficiency (**Figure S2A–2C**). By day 65, neuronal cultures showed evidence of derived maturity, expressing high levels of the post-mitotic neuronal markers MAP2 (microtubule-associated protein 2), NeuN (neuronal-nuclei), and SYP (synaptophysin), as well as specific markers of mature dopaminergic identity, including GIRK2 (G-protein-regulated inward-rectifier potassium channel 2) and TH (tyrosine hydroxylase) (**Figure S2D**).

ZIP14, ZIP8, and ZnT10 all showed relatively high co-expression in mDA neurons with protein expression evident at the cell surface, mitochondria, endoplasmic reticulum (ER), and nucleus (**Figure 1C–1E, S3A-S3C**). Lysosomal expression was not evident (**Figure S3A-S3C**).

In order to investigate the effect of mutant protein on transporter function, inductively coupled plasma-mass spectrometry (ICP-MS) was undertaken to measure intracellular manganese (^55^Mn), iron (^56^Fe), calcium (^44^Ca), and zinc (^66^Zn) levels**, (Figure 1F** and **1G, S3D and S3E**). A disease-specific reduction in intracellular Mn levels was evident in ZIP14 and ZIP8 patient lines, whereas an increase in intracellular Mn levels was detected in ZnT10 lines, under basal conditions. Furthermore, a disease-specific reduction in intracellular Fe levels was also evident in ZIP14 patient lines (**Figure 1G)**. Intracellular levels of Zn and Ca were not disrupted in disease (**Figure S3D and S3E**). Although these metal transporters were originally believed to have a pivotal role in Zn transport,^45,46^ our data provides further evidence that they have a key role in the neuronal transport of Mn. ZIP14 and ZIP8 appear to have a role in Mn import, whilst ZnT10 is a Mn exporter at the cell surface (**Figure 1H** and **1I**). The role of ZnT10 in Mn export was further confirmed by immunoblot analysis of GPP130, a Golgi-specific Mn sensor that is degraded when intracellular Mn levels rise^47^ (**Figure S4A-S4C**).

### Transcriptomic analysis reveals disease-specific dysregulation of key cellular pathways, affecting amongst others, mitochondrial function and calcium signalling, in mDA cultures under basal conditions

In order to determine disease-specific dysregulation of biological processes and cellular pathways, a transcriptomic approach was undertaken with bulk-RNA sequencing.^48,49^ Differentially expressed genes (DEGs) in disease lines were compared to controls, and defined as significant when threshold P-value correction of false discovery rate (FDR) was less than 0.05 and absolute Fold Change (FC) greater than 1.

Disease-specific DEGs were in biological processes related to cellular stress and caspase activation, extracellular matrix (ECM) dysfunction, Ca signalling, and mitochondrial bioenergetics (**Figure S5A–S5C**). Dysregulated subcellular components included the coated vesicular membrane, neuronal projections, plasma membrane, ECM, and organelles including the Golgi apparatus and endoplasmic reticulum (ER) (**Figure S5D–S5I**).

Analysis of Reactome pathways^50^ in ZIP14 patient lines showed that underexpressed DEGs were associated with dysregulation of the cellular response to DNA damage and stress induced senescence, diseases of glycosylation, and organisation/degradation of the ECM (**Figure 2A**). Overexpressed DEGs were mainly associated with pathways linked to collagen biosynthesis, caspase activation, PI3K activation, and neurotransmission (**Figure 2B**). Underexpressed DEGs in ZIP8 patient lines were associated with diseases of glycosylation, signalling pathways (including G protein-coupled receptor “GPCR” ligand binding and caspase activation), peptide hormone synthesis, and transmembrane transport (**Figure 2C**). Transcriptomic defects linked to glycosylation corroborate clinically with patients with SLC39A8-CDG, where Mn-dependent β-1,4-galactosyltransferase is impaired.^23,24^ Overexpressed DEGs were associated with pathways linked to transcription, kinase and phosphatase activity (MAPK activation, IP3/IP4 synthesis, sulphonation) and apoptosis (**Figure 2D**). Underexpressed DEGs in ZnT10 patient lines were associated with ECM organization, cell cycle, diseases of glycosylation, and other pathways linked to transcriptional regulators (**Figure 2E**). Overexpressed genes were associated with pathways linked to diseases of the neuronal system, guanylate cyclase activation through nitric oxide (NO), and neurotransmission (**Figure 2F**).

**Figure 2.**
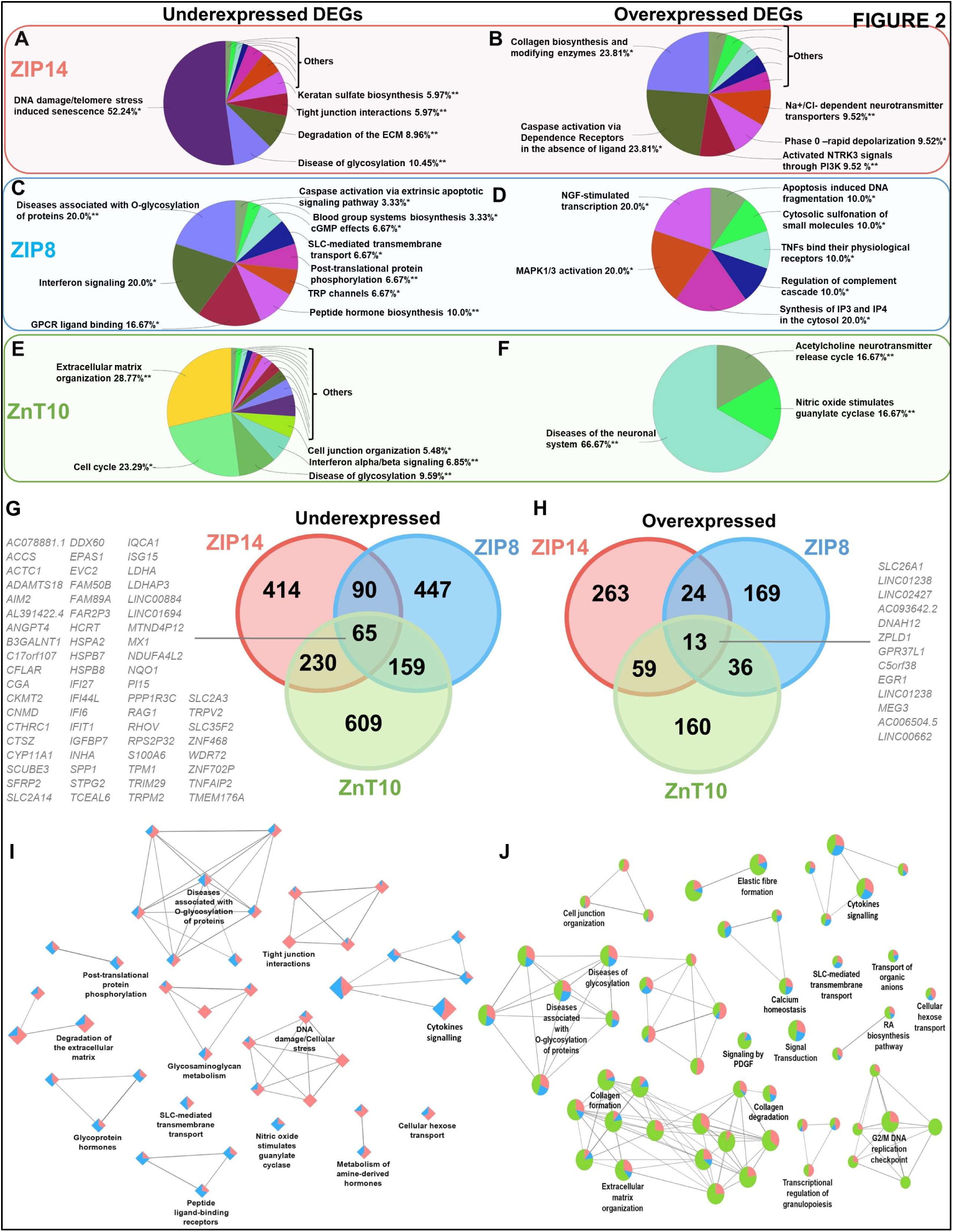
Mn dyshomeostasis leads to dysregulated Reactome pathways in ZIP14, ZIP8, and ZnT10 patient lines under basal conditions, related to Suppl Table S2. **(A-F)** ClueGo analysis of GO terms enrichment for underexpressed and overexpressed DEGs, showing pie chart of Reactome pathways in the ZIP14 (A-B), ZIP8 (C-D), and ZnT10 (E-F) patient lines. Only the GO functional groups exhibiting higher statistically significant differences, using Benjamini-Hochberg P-value correction (FDR<0.05) are shown. **(G-H)** Venn analysis of the shared underexpressed (G) and overexpressed (H) DEGs between ZIP14, ZIP8, and ZnT10 lines. See also **Table S2** for DEGs comparison lists. **(I-J)** ClueGO analysis of GO terms enrichment for Reactome pathways dysregulated in both ZIP14 and ZIP8 patient lines (I) and ZIP14, ZIP8 and ZnT10 lines (J). Network graph nodes represent GO terms and edges indicate shared genes between GO terms. Only the GO functional groups exhibiting higher statistically significant differences, using Benjamini-Hochberg P-value correction (FDR<0.05) are shown.

Of the 2062 DEGs, a total of 78 (65 underexpressed DEGs, 13 overexpressed DEGs) were shared between all three diseases (**Figure 2G** and **2H**). We analysed the DEGs linked to Reactome pathways shared between ZIP14 and ZIP8 lines only, which revealed that Mn deficiency observed in these lines (**Figure 1F**) is linked to shared transcriptomic defects related to pathways of glycosylation and glycosaminoglycan metabolism, DNA damage and cellular stress, peptide ligand-binding receptors, degradation of the ECM, cytokines signalling, and tight junction interactions (**Figure 2I**). Disease-specific dysregulation of Reactome pathways in all three disorders largely comprised genes associated with diseases of glycosylation, collagen formation and degradation, DNA replication checkpoints, signal transduction/cytokine signalling, and calcium homeostasis (**Figure 2J**). Furthermore, for these disorders, PathCards analysis^51^ of shared DEGs highlighted a number of mitochondrial genes involved in glucose and energy metabolism, oxidative phosphorylation, and ATP production, including *SLC2A3*, *LDHA*, *PPP1R3C*, *CYP11A1*, *NQO1*, *NDUFA4L2* and *IGFBP7* (**Table 1**).

### Manganese exposure leads to dysregulation of intracellular metal levels in control and patient-derived mDA cultures

Patients with ZIP14- and ZnT10-related disease have hypermanganesaemia resulting from impaired hepatic Mn homeostasis.^21,52^ Mn overload is also observed in acquired manganism. To recapitulate pathological Mn overload in our cellular model, we exposed control, ZIP14 and ZnT10 patient lines to an acute dose of Mn that did not cause frank cytotoxicity after 48h (**Figure S6A and S6B**).

ICP-MS analysis revealed that acute Mn exposure (100 µM MnCl_2_ for 48h) caused a 40- to 50-fold increase in intracellular Mn levels in all lines, regardless of genotype (**Figure 3A**). Mn exposure was also associated with a marked decrease in intracellular Ca and Fe levels in only ZIP14 patient lines (**Figure 3B** and **3C)**, as similarly observed in other models.^53–55^ Intracellular levels of Zn were not affected in any mDA cultures (**Figure 3D**). Ca and Fe intracellular levels were normal in the ZIP14 isogenic line. Mn exposure also decreased Fe efflux in ZnT10 patient lines, as corroborated by a pulse-chase assay (**Figure S6C-S6F**), and as similarly observed in a SH-SY5Y model of Mn overload.^56^

**Figure 3.**
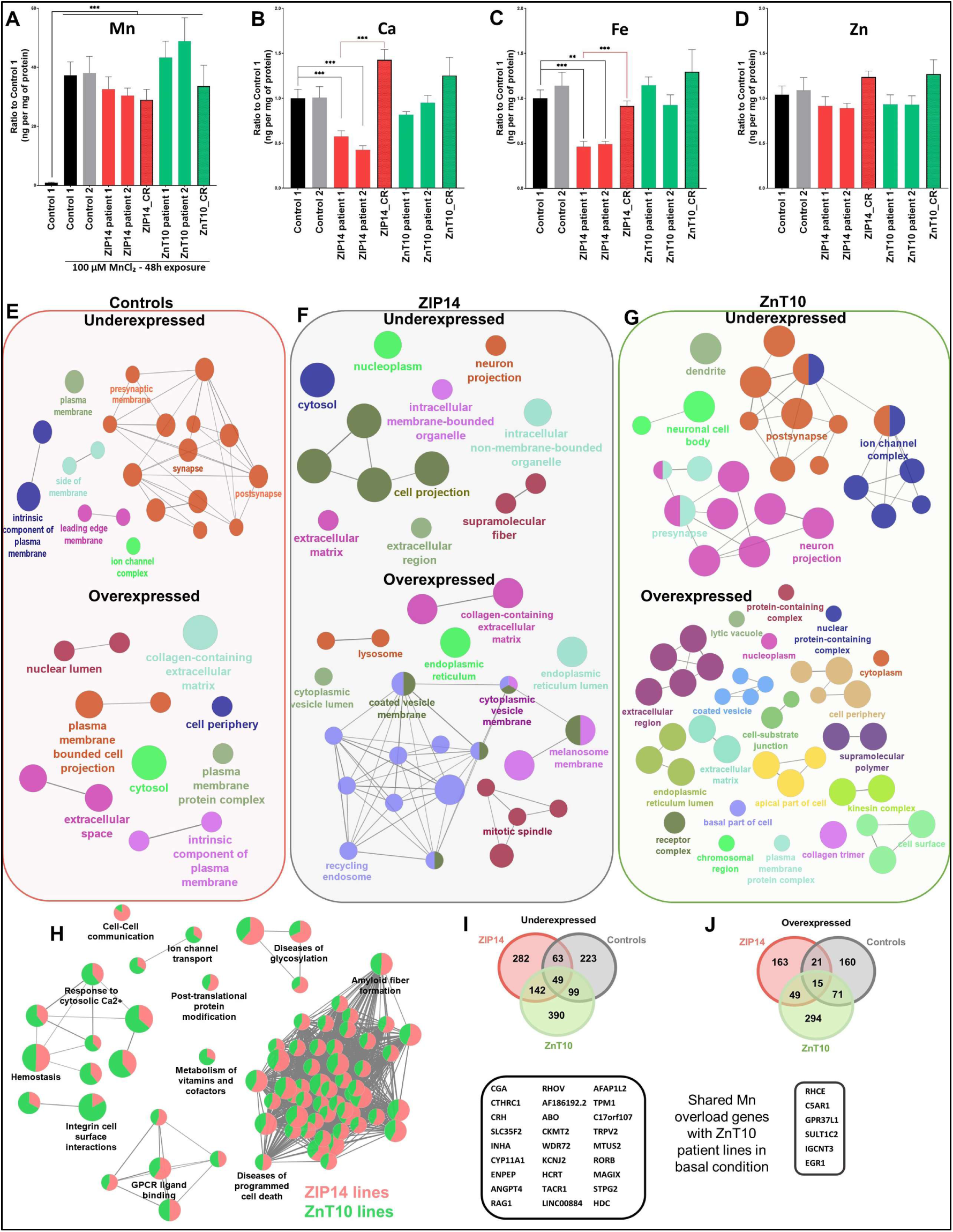
Manganese exposure leads to dysregulated cellular pathways, related to Suppl Table S2. **(A-D)** ICP-MS analysis for intracellular levels of manganese (A), calcium (B), iron (C), and zinc (D) following manganese exposure (100 µM for 48 h). It shows manganese exposure leads to a 40x to 50x increase in intracellular manganese level, as well as affects the intracellular level of both calcium and iron in the ZIP14 patient lines. Zinc levels are unaffected (n=7 to n=13). N= 4 – 15, from minimum 4 independent experiments, ^∗^p=0.05-0.01, **p=0.01-0.001, p***< 0.001, unpaired Student’s t test. Values are given as means ± SEM. **(E-G)** ClueGo analysis of GO terms enrichment for underexpressed and overexpressed DEGs following manganese exposure, showing nodes network of cellular components in controls (E), ZIP14 (F), and ZnT10 (G) lines. Only the GO functional groups exhibiting higher statistically significant differences, using Benjamini-Hochberg P-value correction (FDR<0.05) are shown. **(H)** ClueGO analysis of GO terms enrichment for Reactome pathways dysregulated in both ZIP14 and ZnT10 patient lines exposed to manganese. Network graph nodes represent GO terms and edges indicate shared genes between GO terms. Only the GO functional groups exhibiting higher statistically significant differences, using Benjamini-Hochberg P-value correction (FDR<0.05) are shown. **(I-J)** Venn analysis of the shared underexpressed and overexpressed DEGs between ZIP14, ZnT10, and control lines exposed to manganese. Highlighted genes represent common dysregulated genes also identified in basal conditions for these disorders.

### Transcriptomic analysis reveals enhanced dysregulation of key cellular pathways, including mitochondrial function, calcium signalling and cellular stress in Mn-exposed mDA cultures

Transcriptomic analysis with bulk-RNA sequencing was also undertaken for control and patient lines under Mn exposure. In control lines, analysis of underexpressed DEGs for cellular components showed an association with the synapse (pre- and postsynaptic domains), components of the plasma membrane and ion channel complexes. In contrast, overexpressed DEGs were mainly associated with components of the nuclear lumen, plasma membrane, and cytosol (**Figure 3E**). In ZIP14 patient lines, analysis of underexpressed DEGs showed association with cell projection, intracellular membrane-bounded organelles, neuron projection, parts of the ECM, and supramolecular fibres, whereas overexpressed DEGs were associated with components of the recycling endosome and cytoplasmic vesicle membrane, endoplasmic reticulum, mitotic spindle, and lysosome (**Figure 3F**). In ZnT10 lines, underexpressed DEGs were associated with neuron projection, pre-and postsynaptic processes, ion channel complexes, and dendrites. In contrast, overexpressed DEGs were associated with components of the ECM, cell membrane and its periphery, cytoplasm, ER, and nucleus (**Figure 3G**).

When examining shared dysregulated Reactome pathways in ZIP14 and ZnT10 lines compared to Mn-exposed controls, DEGs related to programmed cell death, amyloid fibre formation, response to cytosolic Ca, diseases of glycosylation, and GPCR ligand binding were identified on Mn exposure (**Figure 3H**). Diseases of programmed cell death and amyloid fibre formation formed the biggest cluster of dysregulated genes, with sub-categories that include defects in transcription, nucleosome assembly, RNA polymerase function, DNA methylation, and oxidative stress, amongst others. Dysregulation of these pathways are consistent with other studies that show Mn exposure can impair the proteasome system, autophagy, and endosomal trafficking, and also cause abnormal amyloid fibre formation.^57^ Moreover, excess Mn is thought to induce exosomal secretion and oligomerisation of α-synuclein, a pathological hallmark of Parkinson’s disease,^58^ which may corroborate the transcriptomic observation of enhanced pathways in the activation of programmed cell death.^29,59,60^

Analysis of shared DEGs between all control and patient lines exposed to Mn identified a total of 64 DEGs (49 underexpressed, 15 overexpressed) shared between all lines (**Figure 3I** and **3J**). Interestingly, 32 of these genes were similarly dysregulated in ZnT10 patient lines in basal condition (27 downregulated and 6 upregulated). This analysis suggests that these genes are the most susceptible targets of Mn overload in this neuronal model, in both basal and Mn-exposed conditions. Pathway analysis categorises these genes mostly to the mitochondria and metabolism (*CYP11A*, *CKMT2*, *HCRT*), calcium signalling (*RHOV*, *TPM1*, *TRPV2*), membrane transport and endocytosis (*WDR72*, *SLC35F2, MAGIX*), neurotransmission (HDC, C17orf107), cellular stress and DNA damage (*EGR1*, *GPR37L1*).

### Manganese dyshomeostasis leads to defects in cellular stress signalling pathways and activated apoptosis

Transcriptomic analysis identified disease-specific DEGs related to cellular stress, MAPK activation, DNA damage and diseases of programmed cell death, in both basal conditions and on Mn exposure (**Figures 2** and **3**). The c-Jun N-terminal Kinases (JNKs)/stress-activated protein kinases (SAPKs) are a subfamily of MAPKs that play a central role in stress signalling pathways involved in gene expression, neuronal plasticity, and regulation of cellular senescence and cell death.^61^ JNK/SAPK pathway activation is postulated to be a key event in many neurodegenerative disorders, including PD, AD, and ALS.^61–63^ As a pathway of interest, we investigated whether the JNK/SAPK pathway was activated in our model, in both basal and Mn-exposed conditions. Immunoblot analysis under basal conditions showed a disease-specific increase in P-JNK, the activated form of JNK (**Figure 4A**). With Mn exposure, although there were no significant differences between control and patient lines (**Figure 4B**), we nevertheless observed an increase in P-JNK expression from basal conditions, which was statistically significant for control and isogenic lines (**Figure 4C**). For patient lines, whilst an increase in P-JNK levels was observed after Mn exposure, it did not reach statistical significance; it is possible that this more modest increase is related to the pre-existing activation of JNK/SAPK in basal conditions.

**Figure 4.**
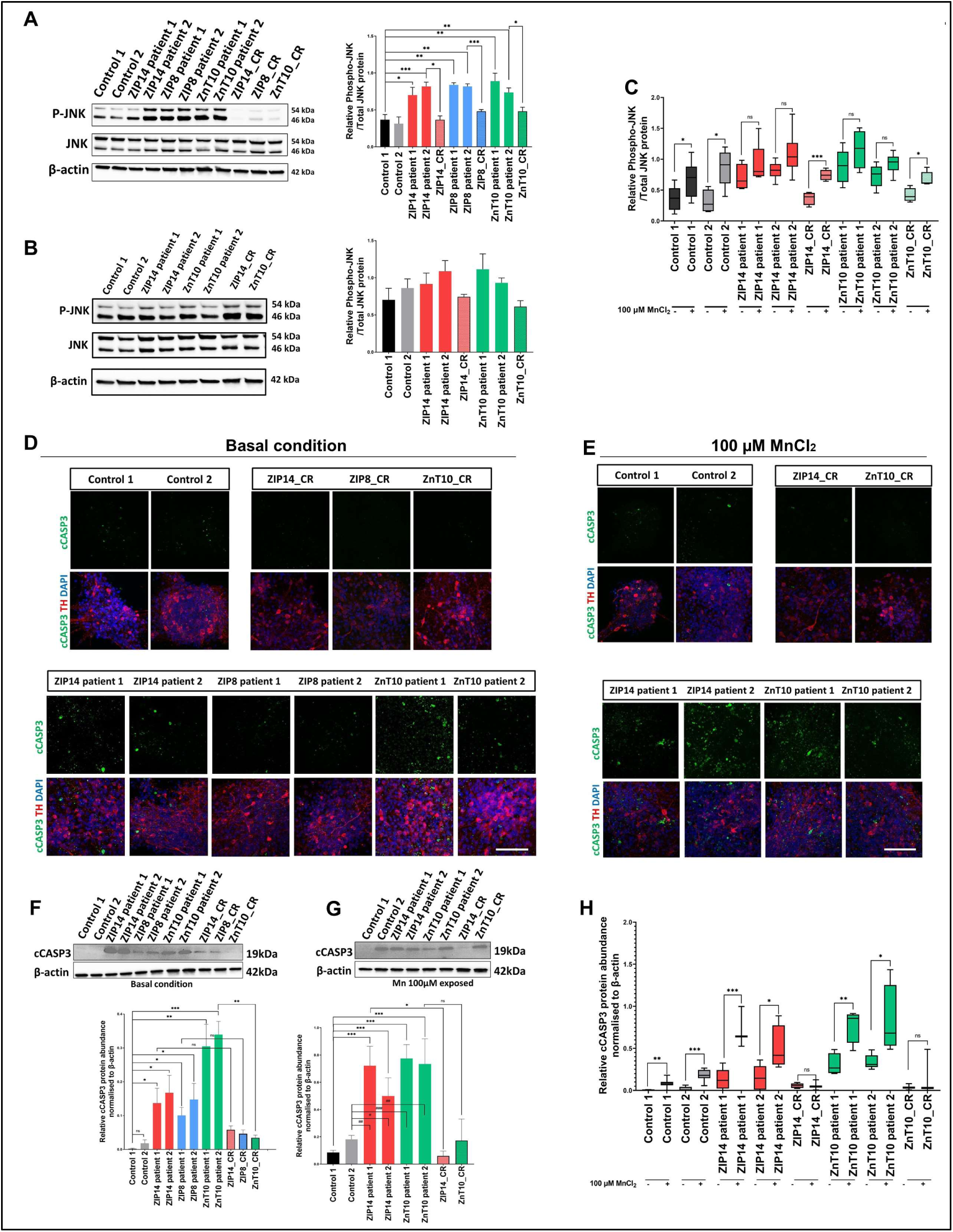
Manganese dyshomeostasis increases cellular stress and markers of early neurodegeneration. **(A-C)** Immunoblot analysis and relative quantification of P-JNK and JNK, in both basal and manganese-exposed conditions. It shows activation of the JNK pathway in basal condition in the patient lines (A), which leads to an increase in JNK activation following manganese exposure (B-C). Box-and-whisker plot shows median with min to max values, n = 3 – 8 from minimum of 3 independent experiments, ^∗^p=0.05-0.01, **p=0.01-0.001, p***< 0.001, unpaired Student’s t test). Values are given as means ± SEM. **(D-E)** Day 65 mDA neurons stained for activated caspase 3 (cCASP3) and Tyrosine hydroxylase (TH) in basal (D) and manganese exposed conditions (E). Scale bar, 100 µm. **(F-G)** Immunoblot and relative quantification for cCASP3 in basal (F) and manganese exposed condition (G). Box-and-whisker plots shows median with min to max values, n = 3 – 11 from minimum of 3 independent experiments, ^∗^p=0.05-0.01, **p=0.01-0.001, p***< 0.001, unpaired Student’s t test. Values are given as means ± SEM. **(H)** Quantification for cCASP3 comparing both basal and manganese exposed conditions. Box- and-whisker plot shows median with min to max values, n = 3 – 11 from minimum of 3 independent experiments, ^∗^p=0.05-0.01, **p=0.01-0.001, p***< 0.001, unpaired Student’s t test. Values are given as means ± SEM.

Activation of JNK/SAPK contributes to the activation of pro-apoptotic events, including expression of pro-apoptotic genes and activation of the caspase cascade in neurons.^64,65^ Our transcriptomic analysis highlighted DEGs in pathways associated with programmed cell death, including caspase activation and apoptosis, both in basal and Mn-exposed conditions (**Figures 2** and **3**). To functionally test the effect of Mn dyshomeostasis on apoptosis-related pathways, we measured the expression of cleaved caspase 3 (cCASP3), a marker for activated apoptosis. Immunofluorescence analysis revealed an increase in cCASP3 in all patient lines under basal conditions, suggesting a disease-specific increase in apoptosis (**Figure 4D**). Following Mn exposure, a further increase in nuclei presenting with cCASP3 staining was observed in both ZIP14 and ZnT10 patient lines when compared to controls (**Figure 4E**). To quantify this, we performed immunoblot analysis for cCASP3 in both basal and Mn-exposed conditions. This analysis showed a disease-specific increase in cCASP3 in basal conditions (**Figure 4F),** and a further increase in cCASP3 in patient lines upon Mn exposure (**Figure 4G**). When comparing both basal and Mn-exposed conditions, we show that Mn appears to induce the activation of cCASP3 in both control and the disease lines (**Figure 4H**). Our results suggest that stress-induced activation of the apoptotic pathway through caspase 3 activation may be an important disease mechanism in the manganese transportopathies.

### Mn dyshomeostasis leads to abnormal mitochondrial bioenergetics in mDA cultures

Mn preferentially accumulates in the mitochondria, where it binds to succinate and malate, directly interfering with mitochondrial respiration and oxidative phosphorylation.^60^ Mn dysregulation is postulated to affect glucose metabolism and ATP production.^5^ Given that transcriptomic analysis identified several DEGs involved in mitochondrial function (**Table 1**), we sought to investigate the effect of Mn dyshomeostasis on mitochondrial bioenergetics in basal conditions and on Mn exposure.

No differences in total mitochondrial mass or mitochondrial DNA content were detected between control and patient lines (**Figure S7A and S7B**). However, disease-specific dysregulation of some components of complex II and IV of the Electron Transfer Chain (ETC) were observed at transcript level (**Figure S7C-S7E**), suggesting that these complexes may be particularly vulnerable to manganese dyshomeostasis. To determine functional impact on the ETC, mitochondrial membrane potential (ΔΨm)-dependent tetramethylrhodamine methyl ester (TMRM) accumulation was measured in both basal and Mn-exposed conditions (**Figure 5A and S7F**). Fluorescence intensity measurements showed a disease-specific reduction in ΔΨm in basal conditions (**Figure S7G**), suggesting that mutations in the ZIP14, ZIP8, and ZnT10 transporters affect mitochondrial membrane potential. We also observed that manganese exposure led to an overall reduction of ΔΨm, regardless of genotype (**Figure 5B**). This suggests that intracellular Mn dyshomeostasis leads to either a disease-specific decrease in ETC complex activity or an increase in proton flux across the inner mitochondrial membrane (IMM), that is worsened on Mn exposure.

**Figure 5.**
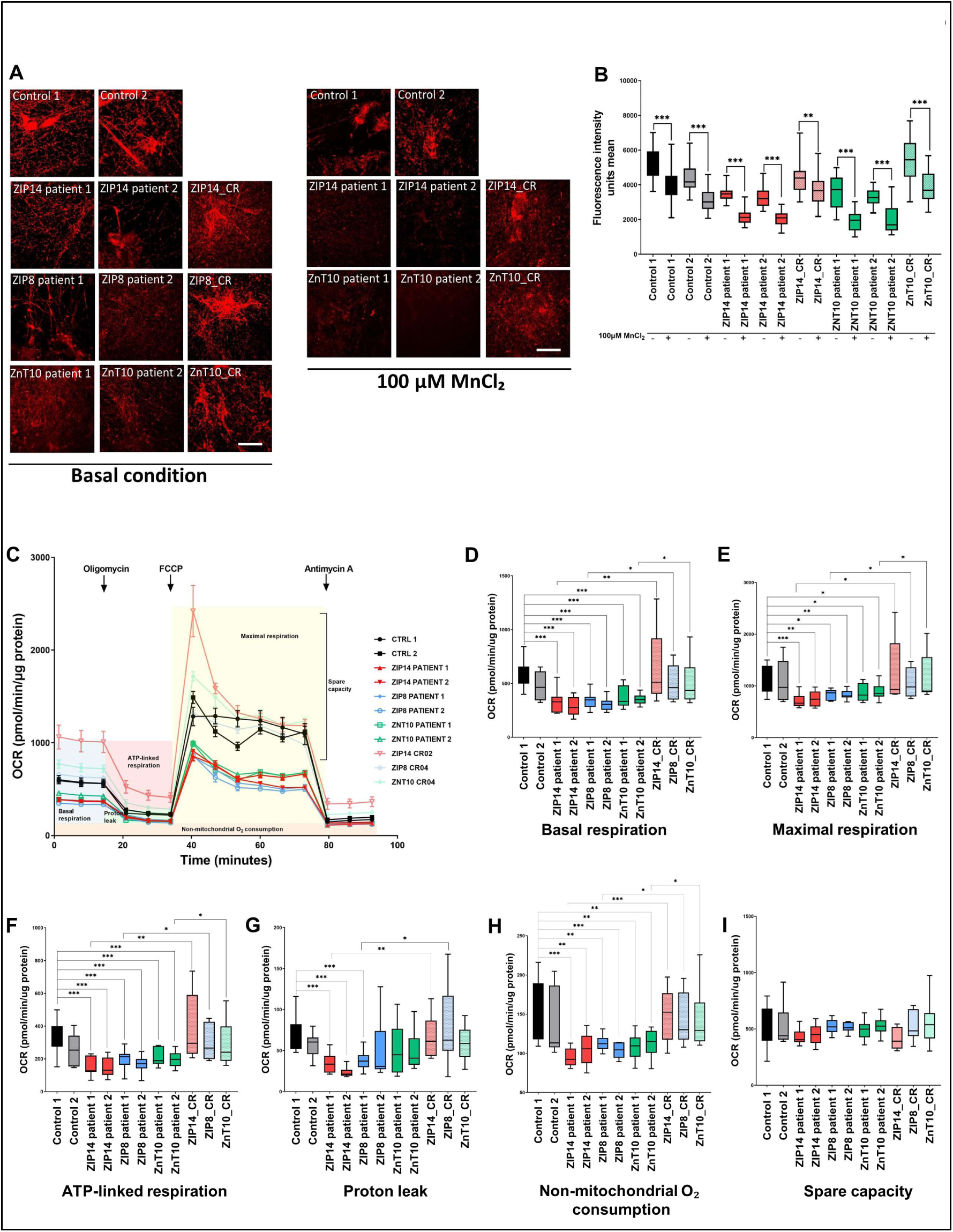
Mn dysregulation leads to defects in mitochondrial bioenergetics under baseline conditions. **(A-B)** TMRM immunostaining in mature neurons, in both physiological and manganese exposed conditions (A). Scale bar = 30 µm. Mean TMRM fluorescence intensity, showing decrease in TMRM intensity following manganese exposure in all lines (B). All lines were independently compared using the two-tailed Student’s t test, ^∗^p=0.05-0.01, **p=0.01-0.001, p***< 0.001 (n=3 to 4, with a minimum of n=6 images/replicate). Box-and-whisker plot shows median with min to max values. Values are given as means ± SEM. **(C)** Representative OCR measurements obtained from Seahorse assays with control and patient neuronal lines normalized to protein content. **(D-I)** Quantification of basal respiration (D), maximal respiration (E), ATP-linked respiration (F), Proton leak (G), non-mitochondrial O2 consumption (H), and spare capacity (I) from Seahorse experiment depicted in (C) with day 65 neurons in basal condition. All lines were independently compared using the two-tailed Student’s t test, ^∗^p=0.05-0.01, **p=0.01-0.001, p***< 0.001. (n>7 biological replicates and n>3 technical replicates). Box-and-whisker plots shows median with min to max values. Values are given as means ± SEM.

To further investigate the effect of Mn dyshomeostasis on ETC function, the oxygen consumption rate (OCR) was measured in mature neurons upon exposure to different stressors of the ETC complexes, in basal conditions (**Figure 5C**).^66^ A significant decrease in basal (**Figure 5D**) and maximal (**Figure 5E**) respiration was observed in all patient lines, suggesting either a decrease in ATP demand, poor ETC integrity, or low substrate availability for all disease lines.^67^ Patient neurons also showed decreased ATP-linked respiration (**Figure 5F**) and reduced proton leak (**Figure 5G**) in ZIP14 and ZIP8 patient 1 neuronal lines, suggesting that there may be overall low ATP demand and ETC damage, at least in some disease lines. Finally, a lower rate of non-mitochondrial O₂ consumption was observed in all patient lines (**Figure 5H**), while spare capacity was unaffected (**Figure 5I**). Overall, our data suggests that Mn dyshomeostasis compromises neuronal mitochondrial bioenergetics across genotypes, with lower energy demand and poor ETC integrity.

We then investigated how Mn exposure affects mitochondrial bioenergetics by measuring OCR in neuronal cultures exposed to 100 µM MnCl₂ (**Figure 6A**). In control lines, we observed a significant decrease in basal, maximal, ATP-linked and non-mitochondrial O₂ consumption upon Mn exposure (**Figure 6B-6E**), suggesting that Mn toxicity may reduce ATP consumption, substrate availability and affect the integrity of the ETC. Proton leak and spare capacity were not affected by Mn exposure (**Figure 6F** and **6G**). OCR was also measured in Mn-exposed ZIP14 and ZnT10 patient lines and compared to their respective isogenic controls, showing a reduction in basal, maximal, and ATP-linked respiration, as well as non-mitochondrial O_2_ consumption between patient and isogenic lines, but not between unexposed and Mn-exposed patient lines (**Figure 6H-6K**). This may suggest that mitochondrial bioenergetics in disease lines are already at a minimal level and toxic exposure to Mn does not cause further reduction of OCR, due to a possible “floor effect”.

**Figure 6.**
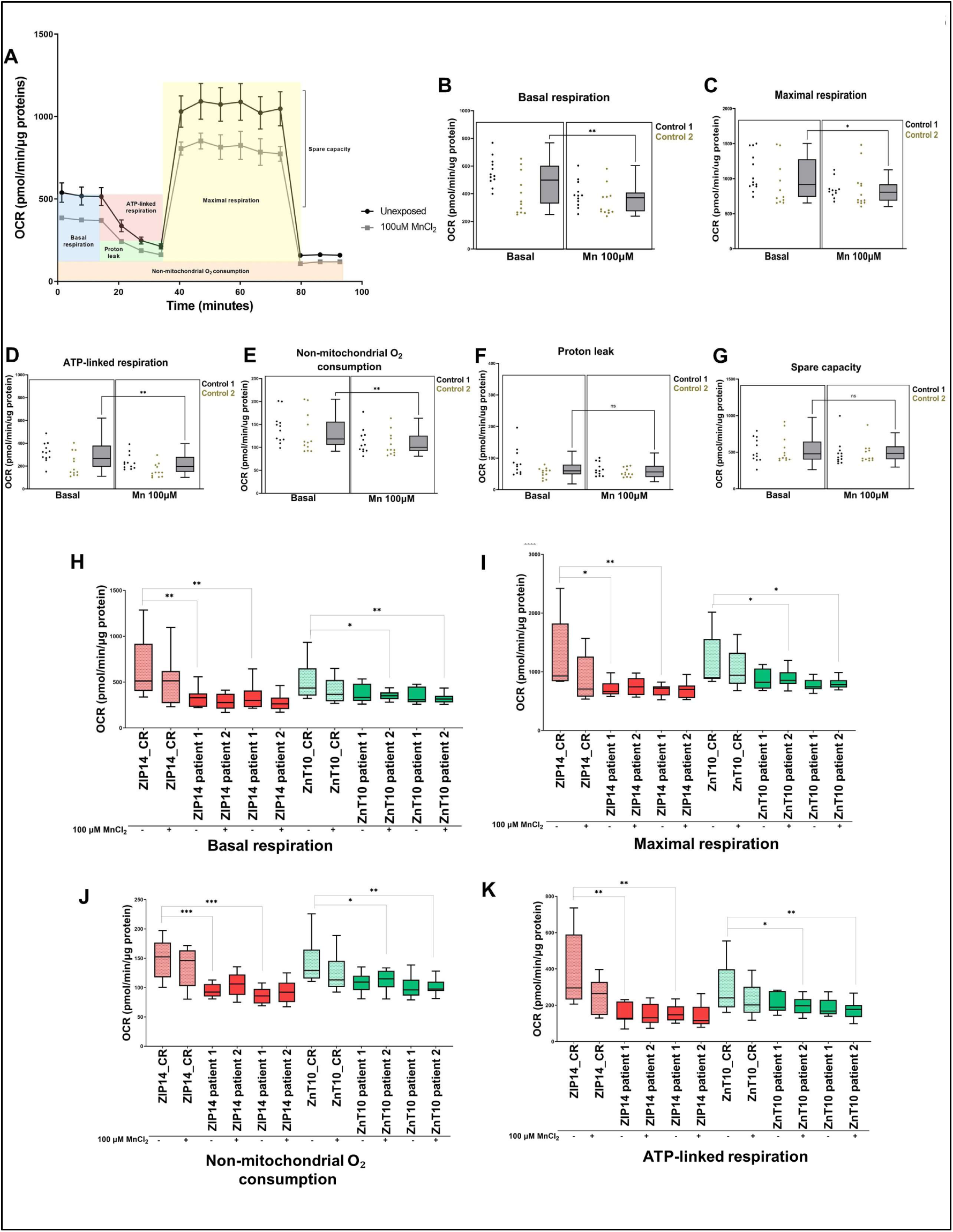
Manganese exposure leads to further defects in mitochondrial bioenergetics. **(A)** Representative OCR measurements obtained from Seahorse assays in control lines normalized to protein content, in manganese exposed condition. **(B-G)** Quantification of basal respiration (B), maximal respiration (C), ATP-linked respiration (D), non-mitochondrial O2 consumption (E), Proton leak (F), and spare capacity (G) from Seahorse experiment depicted in (A) with day 65 control neurons in basal and manganese exposed conditions. Both conditions were compared using the two-tailed Student’s t test, ^∗^p=0.05-0.01, **p=0.01-0.001, p***< 0.001. (n>7 biological replicates and n>3 technical replicates). Box-and-whisker plots show median with min to max values. Values are given as means ± SEM. **(H-K)** Quantification of basal respiration (H), maximal respiration (I), non-mitochondrial O2 consumption (J), and ATP-linked respiration (K) between patient lines and their respective isogenic lines, in both basal and manganese exposed conditions. Conditions were compared using the two-tailed Student’s t test, ^∗^p=0.05-0.01, **p=0.01-0.001, p***< 0.001. (n>7 biological replicates and n>3 technical replicates). Box-and-whisker plots show median with min to max values. Values are given as means ± SEM.

Overall, these analyses suggest that Mn imbalance causes a defect in mitochondrial bioenergetics, by altering the integrity of the ETC. Toxic exposure to Mn also decreases mitochondrial bioenergetics in healthy neurons.

### Mn dyshomeostasis is associated with defects in calcium signalling and endocytosis

Transcriptomic analyses identified a number of DEGs related to Ca^2+^ signalling in states of Mn dyshomeostasis (**Figure 2** and **3**). In mDA neurons, changes in Ca^2+^ homeostasis influence neuronal activity. Mn is postulated to modulate Ca^2+^ channels and regulate neuronal excitability.^68,69^ To determine whether transcriptomic abnormalities associated with Ca^2+^ signalling were functionally evident, control and patient mDA cultures were labelled with CalBryte^TM^ 520 AM, a Ca indicator that allows measurement of Ca signalling events over time. This approach was investigated in basal conditions only, as Mn exposure (100 µM MgCl_2_ over 48h) resulted in dye instability due to Mn quenching. By measuring the ratio between minimal and maximal fluorescence intensity over a 2-minutes period (as a function of ΔF/F0), we observed a disease-specific reduction in the ΔF/F0 fluorescence intensity and frequency of Ca^2+^ transients in patient lines (**Figure 7A and S8A**). To quantify this observed difference, we measured the fluorescence intensity combined with the frequency of Ca^2+^ transients over time (“area under curve”), which showed that Ca^2+^ signalling events are significantly reduced in all patient lines when compared to controls (**Figure 7B**). We treated neurons with Tetrodotoxin (TTX) and NBQX prior to imaging, to confirm that the observed Ca^2+^ fluxes are dependent on action potentials and spontaneous events, respectively. We indeed observed a complete of Ca^2+^ fluxes following TTX treatment, while Ca^2+^ fluxes were mostly conserved following NBQX treatment (**Figure S8B and S8C**).

**Figure 7.**
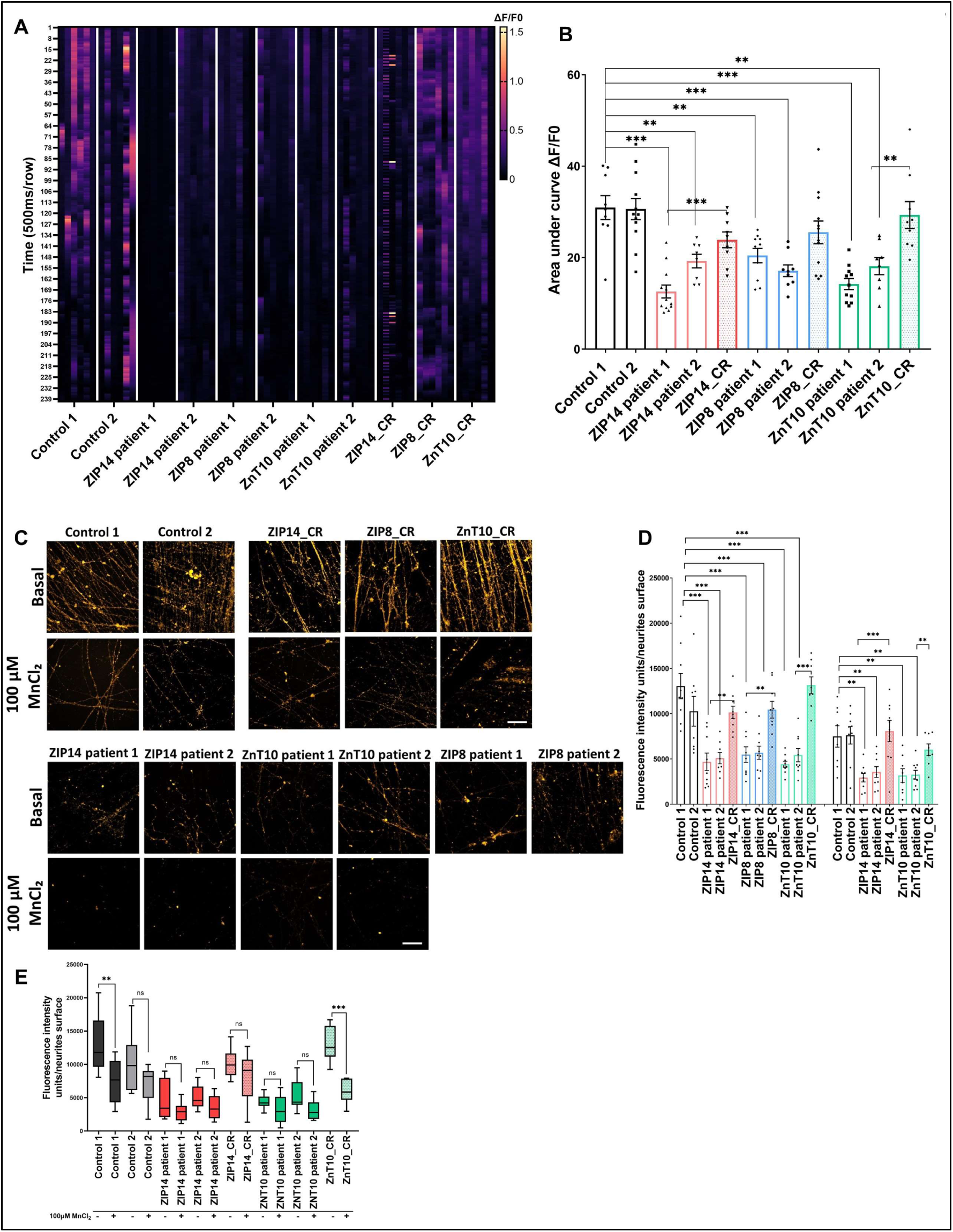
Manganese dyshomeostasis impacts normal calcium signalling and endocytosis. **(A)** Representative figure of the ΔF/F0 fluorescence intensity over a 2 min period, using Calbryte 520 AM calcium indicator, in basal conditions. **(B)** Area under the curve measurements for a minimum of 10 independent cells per images in 3 technical and 3 biological replicates. All lines were independently compared using the two-tailed Student’s t test, ^∗^p=0.05-0.01, **p=0.01-0.001, p***< 0.001. (n>7 biological replicates and n>3 technical replicates). Values are given as means ± SEM. **(C)** Imaging at day 65 with neurons incubated with FM 1-43 dye in both basal and manganese exposed conditions. Scale bar = 20 µm. **(D)** Fluorescence intensity unit measurement for each image (C), normalized to neurite surface. Error bars indicate means ± SEM. All lines were independently compared using the two-tailed Student’s t test for all analysis. (n=9 biological replicates). ^∗^p=0.05-0.01, **p=0.01-0.001, p***< 0.001. (n>7 biological replicates and n>3 technical replicates). Values are given as means ± SEM. **(E) C**omparison of fluorescence intensity units between basal and manganese exposed conditions. Error bars indicate means ± SEM. All lines were independently compared using the two-tailed Student’s t test for all analysis. (n=9 biological replicates). ^∗^p=0.05- 0.01, **p=0.01-0.001, p***< 0.001. (n>7 biological replicates and n>3 technical replicates). Box-and-whisker plot shows median with min to max values. Values are given as means ± SEM.

Ca^2+^ modulates axon outgrowth, neuronal survival, synaptic strength, and neurotransmission, by triggering neurotransmitter release.^70^ Vesicular endocytosis and exocytosis, coupled with Ca^2+^ signalling, are key processes in normal neurotransmission.^71,72^ Importantly, transcriptomic analysis indicated endocytosis as a pathway of interest, with DEGs in both disease lines and on Mn exposure including dysregulation of *SYT2*, *CaM*, *SYNPO*, *SNCA*, and *SYP* amongst others (**Table 1 and Table S2**).

To validate our transcriptomic data, we sought to determine whether endocytosis was disrupted by Mn dyshomeostasis in our neuronal model. Vesicle recycling was investigated through an established FM 1-43 uptake assay. In active neurons, this dye is internalised within recycled synaptic vesicles, staining the nerve terminals and internal membranes.^73^ Control and patient mDA cultures were incubated with FM 1-43 in basal and Mn-exposed conditions (**Figure 7C**). Disease-specific reduction of staining intensity, indicative of impaired FM 1-43 uptake, was evident in basal conditions, which was further decreased in all lines upon Mn exposure (**Figure 7C**). Quantification, by measurement of the ratio of fluorescence intensity units per neurite surface, confirmed a statistically significant reduction of fluorescence intensity in the disease lines in basal conditions, which was further decreased for controls, ZIP14 and ZnT10 patient lines on Mn exposure (**Figure 7D).** The additional decrease in fluorescence intensity on Mn exposure did not reach statistical significance for patient-derived lines, which may suggest that endocytosis is already significantly impaired in basal conditions (**Figure 7E**). Mn exposure also leads to a reduction in FM 1-43 fluorescence intensity in some control lines. Overall, this analysis implies that both acquired and inherited forms of Mn dyshomeostasis negatively impact neuronal endocytosis.

## Discussion

Mn is a trace metal that is essential to life. Although defective Mn homeostasis is implicated in a broad range of human diseases, the underlying pathophysiology of Mn dyshomeostasis is poorly understood. To better understand the role of Mn in the human brain, we have generated a patient-derived mDA neuronal system to study the molecular consequences of both acquired and inherited forms of Mn dyshomeostasis. Using this neuronal system, we were able to model both systemic and intraneuronal Mn deficiency as well as states of Mn overload and toxicity. Indeed, our study simulates the systemic and intraneuronal Mn deficiency observed in SLC39A8-CDG disease,^23,24,26^ and the systemic overload observed in acquired manganism and in the inherited HMNDYT1 & 2 disorders.^11,19,21,39,74^ Our model also sheds further light on the complex downstream effects of SLC39A14 mutations: whilst deficiency of this gene leads to systemic Mn accumulation in HMNDYT2 patients (through loss of hepatic uptake), we do not observe Mn overload in our neuronal model unless the system is exposed to Mn. The complex function of SLC39A14 as a Mn importer manifests differently in the brain, or at least in DA neurons, compared to other organs. Our SLC39A14 model therefore offers the added advantage of investigating the potential effects of intraneuronal Mn deficiency. Our findings align with the partial Mn deficiency seen in a *slc39a14*-KO zebrafish model,^53^ suggesting involvement of diverse cellular and organ-specific mechanisms for Mn homeostasis. Overall, our model reveals that both Mn overload and deficiency result in a number of common dysregulated pathways, including those governing mitochondrial function, cellular stress, neurodegeneration, endocytosis and Ca^2+^ signalling. Our model is the first humanised neuronal model of Mn-related disease and confers several advantages when compared to existing animal and cellular models, that are often limited by species-specific differences from the human brain.^29^

Our mDA model confirms that ZIP14, ZIP18 and ZnT10 have fundamental roles in the neuronal cellular import (ZIP14, ZIP8) and export (ZnT10) of Mn. In disease states, deficiency of these transporters in the gut and enterohepatic system leads to dysregulation of blood Mn levels. Murine models for these disorders confirm the crucial role for these Mn transporters in the liver and intestine.^75–78^ Whilst systemic hypo- or hypermanganesaemia are considered to be major factors in disease causation, the primarily neurologic phenotype argues for a prominent role of the disruption of intracellular Mn levels in the brain in the pathogenesis of these conditions. As illustrated by our transcriptomic analysis, defective Mn homeostasis in neurons is associated with important sequelae across a broad range of molecular and cellular process, related to glycosylation, ECM, collagen, endocytosis, calcium signalling, glucose and energy metabolism, cellular stress and apoptosis - all of which are critical to normal neuronal function.

Our iPSC-derived neuronal model revealed that Mn dyshomeostasis is linked to activation of the stress-responsive JNK/SAPK signalling pathway and initiation of the apoptosis cascade, through caspase 3 activation. An increase in cCASP3 observed in our *in vitro* system may herald future neurodegeneration. Reports suggest that chronic exposure to Mn increases the risk of developing neurodegenerative diseases, including AD, PD, ALS, and HD.^16,79^ Post-mortem studies have shown neuronal damage in the globus pallidus and striatum, with sparing of the SNc and absence of Lewy bodies.^80^ Whilst autopsy data is scant for the monogenic Mn transportopathies, studies to date reveal significant neuronal loss within the basal ganglia, in particular the globus pallidus, dentate nucleus, and cerebellum, as well as vacuolisation and myelin loss throughout the white matter in one case of *SLC39A14* deficiency ^21^. In a *SLC30A10* patient, post-mortem analysis showed neuronal loss, astrocytosis, myelin loss, and spongiosis in the basal ganglia.^81^ The identification of early neurodegenerative markers in a humanised cell model is therefore important, especially given that neurodegenerative phenotypes are not always recapitulated in animal models. *Slc39a14* knockout mice do not show mDA neurodegeneration, even when aged.^82^ Similarly, a pan-neuronal/glial *Slc30a10* knockout mice model did not show signs of dopaminergic neurodegeneration, even in adult mice.^83^ Conversely, a study in a *SLC30A10*-mutant model of *Caenorhabditis elegans* showed dopaminergic degeneration following Mn exposure.^84^ Such discrepancies between animal models therefore highlights the need for more advanced humanised systems to better understand the pathophysiological effects of Mn in the brain.

The intricate interplay between Mn dyshomeostasis and mitochondrial function has emerged as a central theme in our model, with dysregulation of mitochondrial genes within the electron transport chain (particularly for complexes II and IV), abnormal MMP, disruption of ATP-linked respiration and non-mitochondrial O_2_ consumption, and reduction in cellular energetic demands and response to metabolic fluctuations. Our study builds upon established findings in the field, corroborating the presence of mitochondrial defects in states of Mn dyshomeostasis.^85–89^ These studies collectively showed the deleterious effects of excessive Mn on mitochondrial respiration, ROS production, and ATP production, amongst others ^4^. Our findings together emphasize on the importance of Mn normostasis in mitochondrial bioenergetics. The effect of Mn dyshomeostasis on mitochondrial function may involve many mechanisms, including dysfunction of the TCA cycle, lack of enzyme intermediates, accumulation of toxic substances, and ROS production. For instance, Mn preferentially accumulates in the mitochondria and acts as an activator of the gluconeogenesis enzymes phosphoenolpyruvate carboxykinase and pyruvate carboxylase.^5^ In addition, isocitrate dehydrogenase is a Mn-dependent and the rate-limiting enzyme of the Krebs cycle, which converts isocitrate to α-ketoglutarate with production of CO_2_ and NADH.^5^ Dysregulated enzymatic activity due to Mn dyshomeostasis may therefore contribute to abnormal mitochondrial respiration. Elevated Mn is also known to bind to TCA intermediates succinate, malate, and glutamate, directly interfering with mitochondrial respiration and affecting oxidative phosphorylation.^4^ Mn is involved in the mitochondrial oxidative stress response and ROS detoxification by acting as a cofactor for the Mn-dependent superoxide dismutase 2 (SOD2). Dysregulation of SOD2 levels can result in an imbalance between its antioxidant function and pro-oxidative potential.^90^

Our model also sheds light on the role of Mn in neuronal Ca^2+^ signalling and spontaneous firing activity in mDA neurons. Ca^2+^ signalling is essential for the physiological function of healthy neurons and perturbed Ca^2+^ homeostasis has been implicated in the pathogenicity of several neurodegenerative disorders, including AD, PD, ALS, and HD.^91^ In dopaminergic neurons, changes in Ca^2+^ homeostasis influence neuronal activity, and Mn has been shown to modulate Ca^2+^ channels, therefore regulating dopaminergic neuron excitability.^68,69^ Evidence from the literature suggest that some store-operated Ca^2+^ channels are permeable to Mn, representing a mechanism for Mn influx into the CNS.^3,91^ The competitive nature of Mn with Ca^2+^ ^92^ may interfere with Ca^2+^ intracellular intake and signalling, which could potentially explain the Ca^2+^ signalling defects observed in our neuronal model. Evidence from the literature also suggests that mitochondria play a key role in Ca^2+^ homeostasis, regulating the absorption and storage of Ca^2+^ ions.^93^ In an iPSC-derived mDA model for PD, mitochondrial failure was shown to cause calcium dyshomeostasis which led to synaptic dysfunction of dopamine neurons.^94^ It is therefore plausible that the observed defects in Ca^2+^ signalling within our neuronal model may also partly be explained by the observed mitochondrial dysfunction in our patient lines, which warrants for future investigation.

It is also apparent that Mn dyshomeostasis leads to defects in endocytosis in the iPSC-derived model, which, to our knowledge has not been directly reported in other Mn-related disease models. Normal Ca^2+^ signaling is of paramount importance for triggering endocytosis and exocytosis.^70,71^ Evidence indeed suggests that intracellular Ca^2+^ regulates both exocytosis and endocytosis, via different proteins, including synaptotagmin, calmodulin, and calcineurin as well as Ca^2+^ micro-domain.^71^ This supports the link between Ca^2+^ signaling and endocytosis, where an increase in Ca^2+^ signaling events increases the rate of endocytosis, and a decrease in Ca^2+^ signaling reduces the endocytosis rate.^95^ It is therefore possible that the observed endocytosis defects could be attributed to the abnormal calcium signaling observed in our neuronal model. Additionally, the release of neurotransmitters in the brain relies on the processes of endocytosis and exocytosis, which is crucial for the health and normal functioning of neurons. Studies in *Slc39a14*-KO and pan-neuronal/glial *Slc30a10*-KO mice have demonstrated deficiencies in dopamine (DA) release within the striatum, likely attributed to impairments in endocytosis and exocytosis due to Mn overload.^82,83^ Moreover, analysis of gene expression in *Slc30a10*-KO mice has revealed abnormalities in genes governing synaptic transmission and neurotransmitter function,^96^ while both pre- and post-synaptic gene defects, along with disruptions in Ca^2+^ signaling, were observed in the *slc39a14*-KO zebrafish model.^53^ The identification of shared DEGs across species, including in our human models, further strengthens the potential link between Mn homeostasis and normal neurotransmission.

The clinical treatment of both inherited and acquired disorders of Mn dyshomeostasis remains highly challenging. For many patients, the mainstay of therapy is largely symptomatic, with palliative measures for symptom control using chelation therapy (to remove Mn) and levodopa administration (to increase DA production and treat the movement disorder).^29^ Both therapies are controversial as while they provide limited benefit for some patients, they are ineffective in others who continue to deteriorate over time.^22,97–99^ There is also an additional burden with conventional chelation therapies, as they require regular intravenous injections and patients must be closely monitored to avoid potential adverse effects. Current chelation therapies are also not specific to Mn (leading to off-target effects such as anaemia, low levels of magnesium and potassium, low blood pressure) and their inability to cross the blood-brain barrier (BBB) make them a poor option to treat toxic accumulation of brain Mn.^21,100,101^ Importantly, our model suggests that Mn dyshomeostasis results in multifaceted neuronal pathophysiology and as such, targeting a singular dysregulated pathway is unlikely to be disease-modifying or curative.^29,100,102^ Similarly, the detrimental effect of excessive Mn on our model highlights the need of dose optimization and careful therapy control in the context of Mn supplementation in SLC39A8-CDG.^27,103^ Potential future options could include the development of more effective, Mn-specific chelators that easily cross the BBB. For the monogenic Mn transportopathies, genetic therapies have real potential to treat the underlying loss-of-function defects in ZIP8, ZIP14 and ZnT10 deficiency. This could be through RNA therapeutics or AAV-mediated DNA therapies. Recent advances and successes in gene therapy using adeno-associated viral (AAV) vectors for gene delivery in inherited neurological disorders provide great hope for the applicability of such approaches to the inherited monogenic Mn transportopathies.^104^

Specifically, targeted AAV8 therapy aimed at the liver increased the expression of *SLC30A10* and aided in mitigating systemic Mn overload in *Slc30a10*-KO mice.^105^ Rectifying SLC30A10 deficiency in the liver was adequate to alleviate disease severity, but it did not completely restore Mn levels to those of wildtype mice. Addressing tissue specificity is therefore imperative for more precise targeting and enhanced efficacy and it may be that both systemic and brain delivery of gene therapy may be required for effective rescue. Nonetheless, these AAV-based methods described provide important first proof-of-concept, and hold great promise as potential treatment options for patients with HMNDYT1 & 2, and SLC39A8-CDG. The advent of CRISPR-Cas9 technology and gene editing capabilities will also likely provide future therapeutic avenues.^106–108^ Nevertheless, any form of genetic therapy poses significant translational challenges; refining technical modalities and delivery systems to maximise safety and optimise efficacy within a crucial therapeutic window before irreversible brain damage will be pivotal for successful implementation in patients.^29^

In conclusion, development of a humanised iPSC-derived neuronal model of Mn dysregulation has provided important insights into the crucial role of Mn in normal neuronal function. Mn is both an essential micronutrient and potential neurotoxicant,^42^ and tight regulation of intracellular and extracellular Mn is therefore key to prevent perturbation of a broad range of molecular processes in the brain. Improved understanding of these process will not only have implications for both rare and common neurodevelopmental and neurodegenerative diseases, but also for occupational and environmental settings, informing public health policy and environmental safety.

## Acknowledgements

We sincerely thank our patients and their families for participating in this study. We thank Dr. Gleeson (Gleeson lab, University of California in San Diego, USA) for isolating the ZnT10 patient fibroblasts from skin biopsies provided by Dr. Zaki (National Research Centre, Cairo, Egypt); Dr Derek Burke (Great Ormond Street Hospital for Children NHS Foundation Trust, Department of Chemical Pathology Enzyme unit, London, UK) for isolating the ZIP14 patients and ZnT10 patient 1 fibroblasts from skin biopsies; Dr Karin Tuschl and Dr Peter Clayton for coordinating with patients for fibroblast collection and consent; Dr Philippa Mills for supervising Dr. Dimitri Budinger’s PhD thesis work included in this manuscript. We thank Dr. Ningning Zao (University of Arizona, USA) for providing the ZIP14 antibody. We are grateful to Dr. Olivia Gilham and Dr. Preethi Sheshadri for Seahorse introduction; Dr Nandaki Keshavan and Dr. Shamima Rahman for their expertise in mitochondrial OXPHOS activity.

## Materials and methods

### Resources table

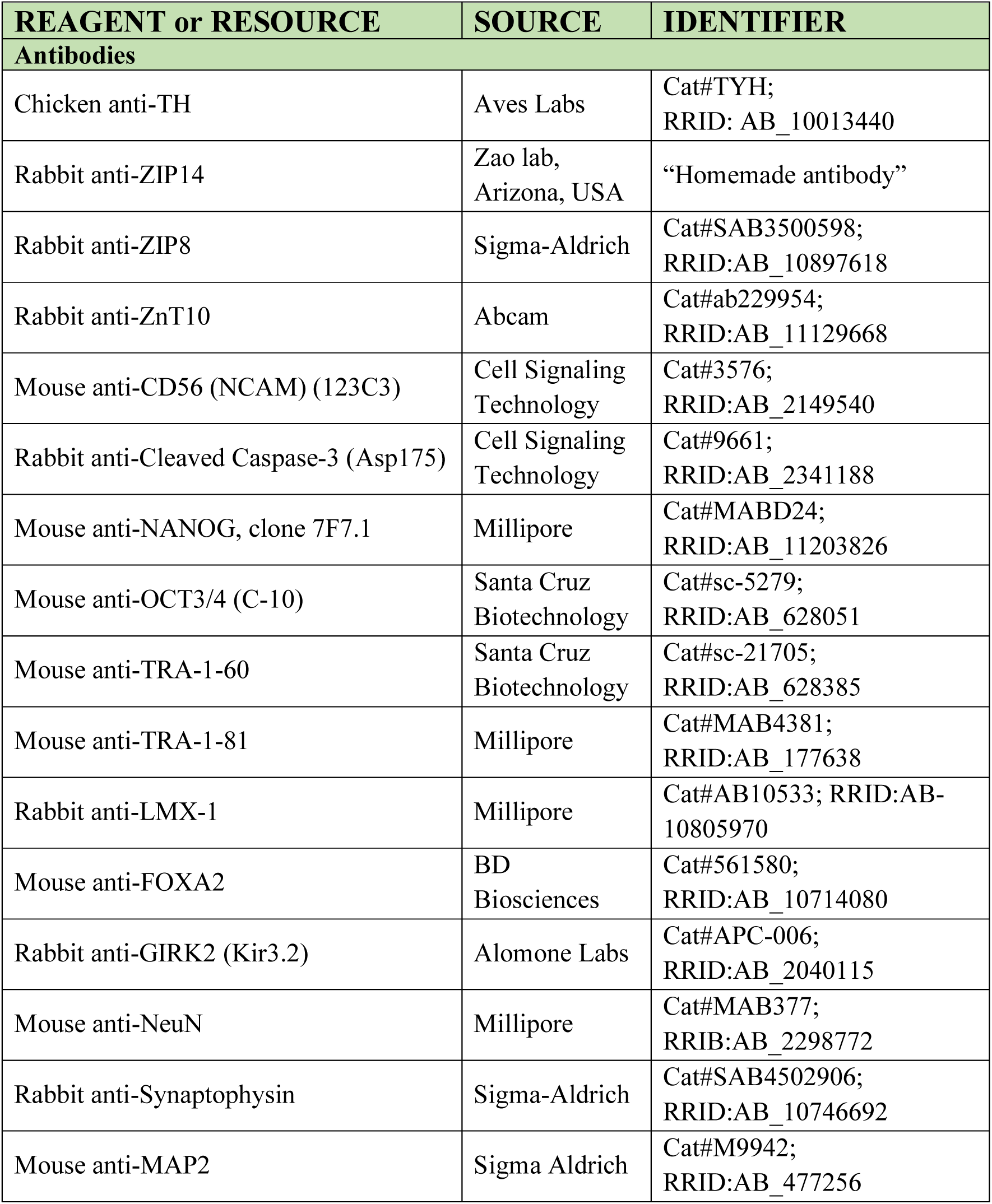
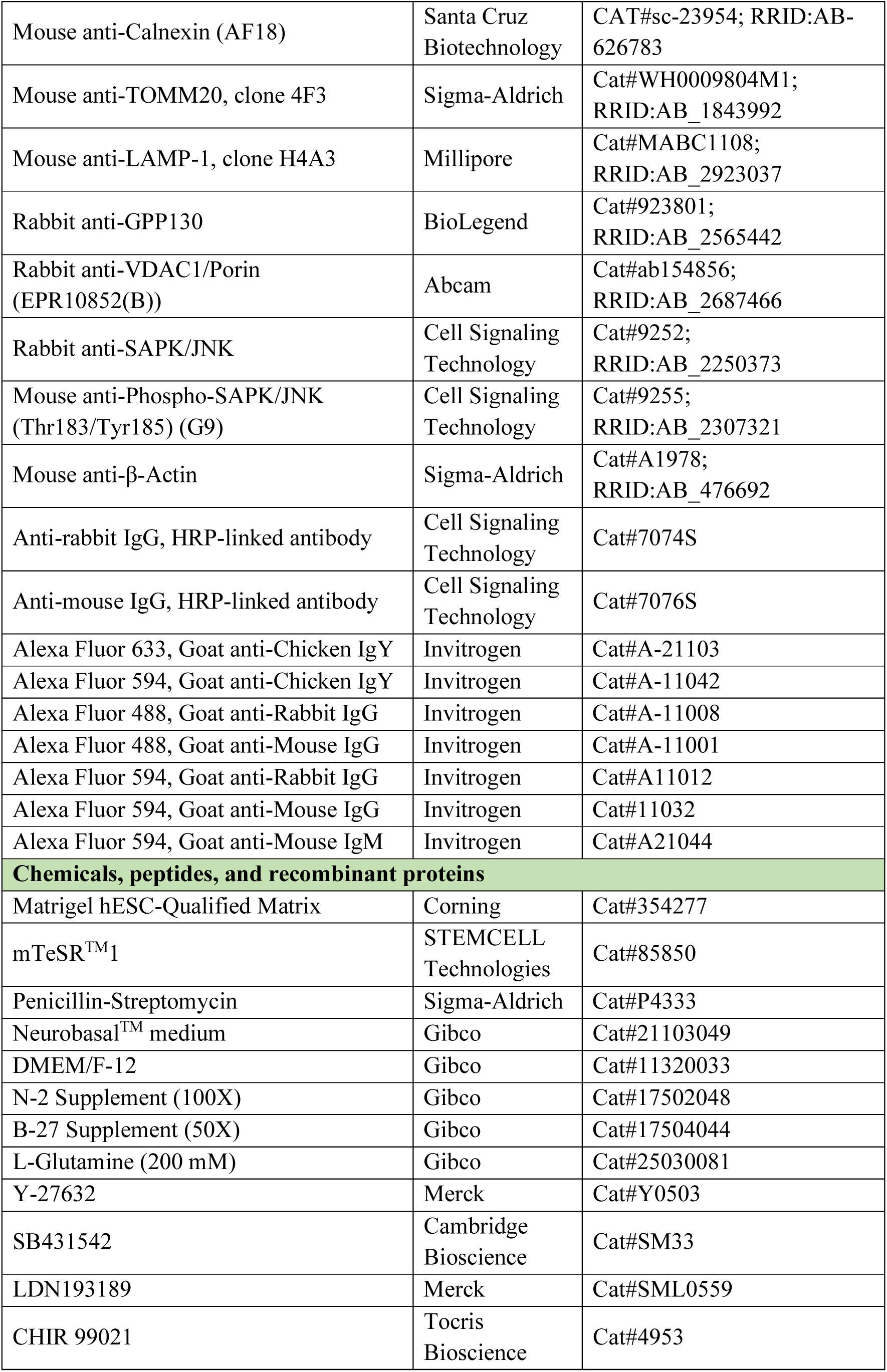
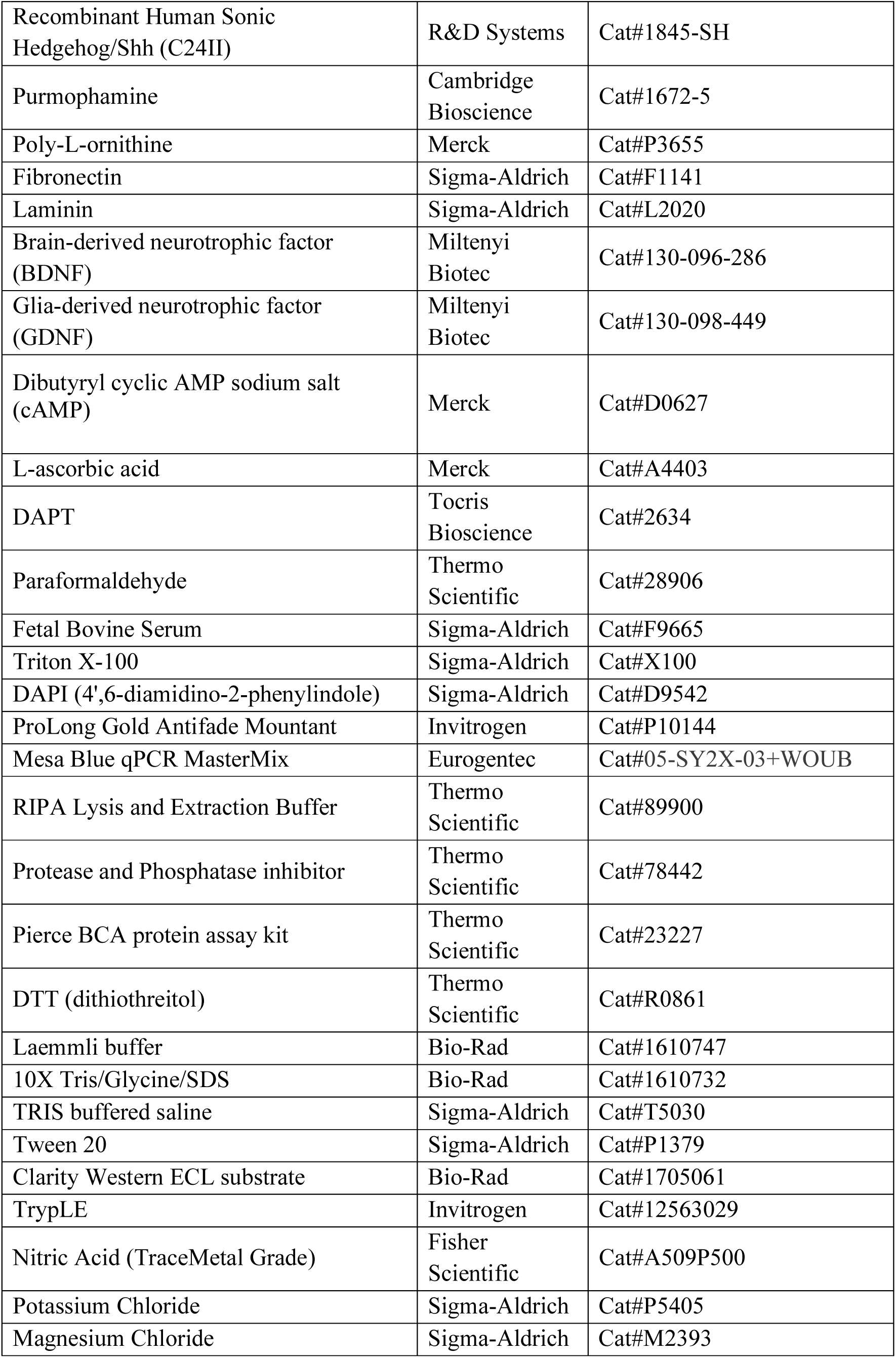
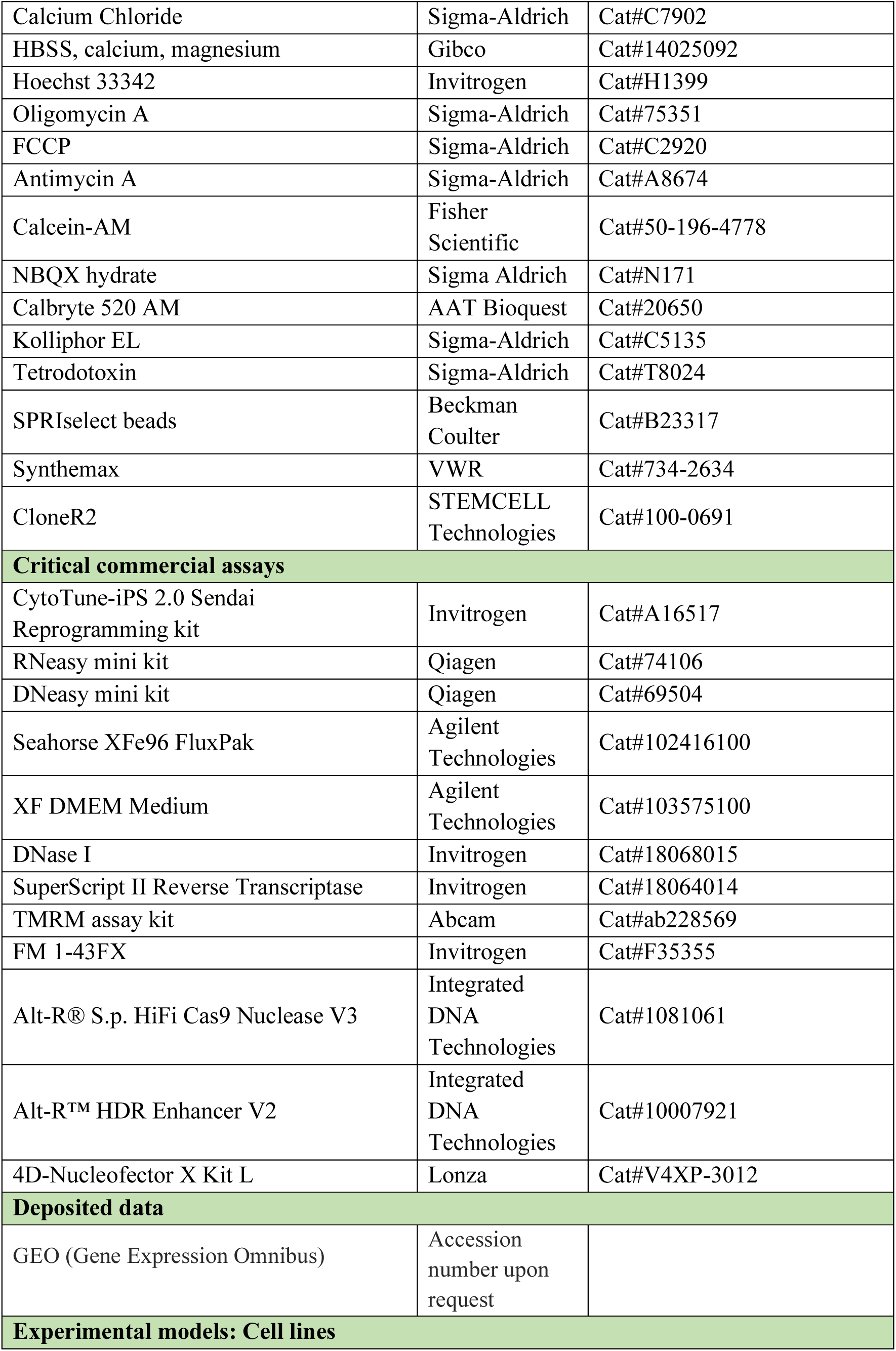
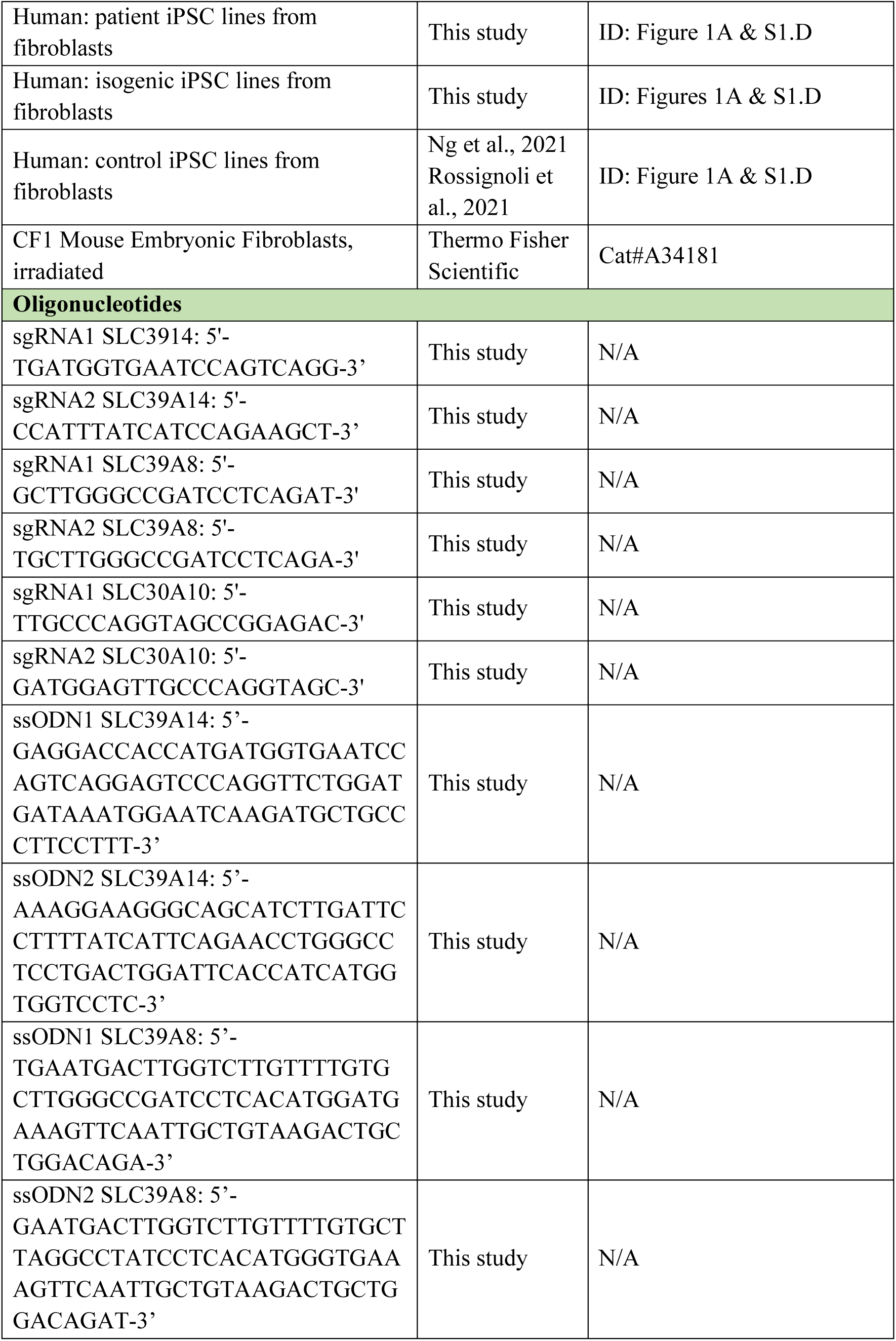
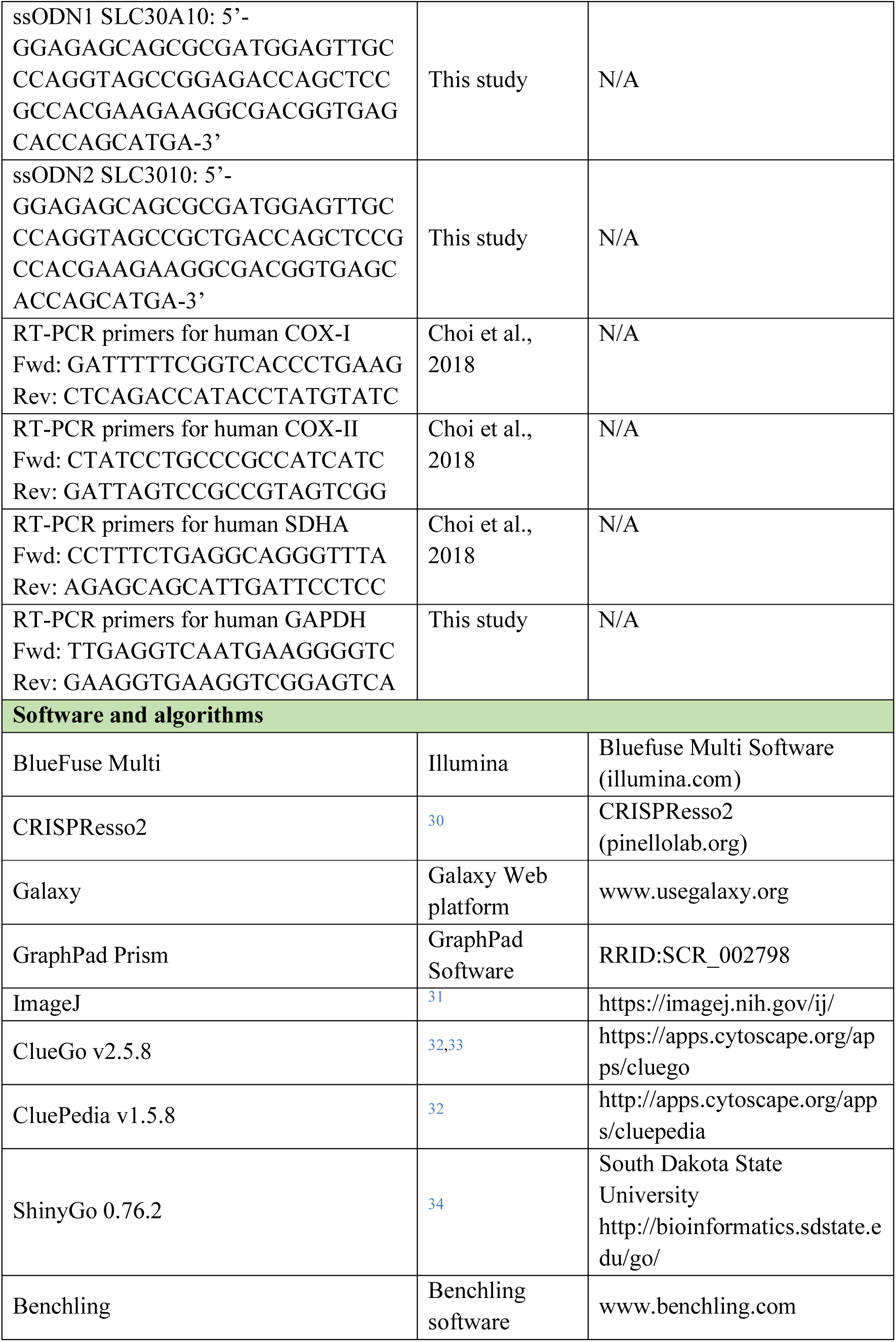

### Induced pluripotent stem cells generation, characterisation and maintenance

Isolation of human dermal fibroblasts (HDFs) from skin biopsies was performed by the Enzyme Unit, Chemical Pathology, Camelia Botnar’s Laboratories, Great Ormond Street Hospital, London, UK and by the Gleeson Laboratory for Paediatric Brain Disease, Rady Children’s Institute for Genomic Medicine, University of California, San Diego, USA. Written informed consent was obtained from all participants (for the SLC39A14 patient 1 and 2 and SLC30A10 patient 1; Great Ormond Street Hospital patients; REC reference 13/LO/0171 and 13/LO/0168, for the SLC39A8 patient 1; Western Sydney Genetics Program consent HREC/10/CHW/114, for the SLC39A8 patient 2; Ethik-Kommission Westfalen-Lippe 2012-373-f-S, for the SLC30A10 patient 2; STDF-33650-Ethical approval 20105). The protocol used for generation of hiPSCs is adapted from the CytoTuneTM-iPS 2.0 Sendai Reprogramming Kit (Invitrogen), as described previously.^35,36^ Generated iPS cells were cultivated on Matrigel-coated plates (Corning) and mTeSR^TM^1 complete medium (StemCell Technologies) supplemented with 1 % P/S. Genome integrity was assessed by the Illumina Human CytoSNP- 12 v2.1 beadchip array and by Sanger sequencing of patient’s mutations prior to, and after reprogramming. Pluripotency of the lines was assessed by Epi-Pluri-Score (Cygenia, Epigenetic Diagnostics, Aachen, Germany), and by immunocytochemistry for pluripotency markers OCT4 (1:50; Santa Cruz), NANOG (1:500; Millipore), TRA-1-60 (1:200; Santa Cruz), TRA-1-81 (1:200; Millipore). Sendai virus clearance, PCR detection for pluripotency markers and spontaneous *in vitro* differentiation ‘STEMdiff^TM^ Trilineage Differentiation Kit (StemCell Technologies) were also performed (data not shown). Two clones per iPSC line were originally generated and characterised. One clone per line was then carried forward for analysis.

### CRISPR/Cas9 genome editing

The protocol used for the generation of isogenic control lines is based on a published protocol.^37^ In brief, two independent single guide RNA (sgRNAs) were designed to target the following mutations in SLC39A14 patient 1, c.[1407C>G] (NM_001128431.4), in SLC39A8 patient 1, c.[338G>C] (NM_001135146.2) and in SLC30A10 patient 2, c.[77T>C] (NM_018713.2).

Single-stranded DNA oligonucleotides (ssODN) were designed to comprise a silent mutation either in the PAM sequences or elsewhere in the ssODN. For each isogenic line, a total of 8 x 10^5^ cells were electroporated together with the pre-assembled Cas9-sgRNA complex and respective ssODN. Electroporated cells were transferred in pre-warmed mTeSR medium with 10 % CloneR2 (StemCell Technologies) and HDR enhancer (IDT, 30 µM). Cells were incubated at 32°C/5 % CO_2_ for 48 h. Medium was changed 16 h post electroporation with pre-warm mTeSR medium with 10 % CloneR2. After 48 h, cells were incubated at 37°C/5 % CO_2_, with daily mTeSR medium change until they reached 70-80 % confluency. Low density was performed to achieve single cell plating for colonies picking. Individual round colonies were picked and duplicated for both genotyping and freezing purposes. Genotyping of individual clones was assessed following a 3 PCR steps, as follows: i) 500 bp PCR around edit, ii) nested 200 pb PCR with specific appends, iii) barcoded PCR with i5 Index and i7 Barcoded MiSeq primers. MiSeq sequencing was performed in collaboration with UCL Genomics, Zayed Centre for Research into Rare Disease in Children, London, UK. MiSeq output (fastq files) were loaded into CRISPResso2 for data analysis.^30^

### Midbrain dopaminergic neuronal differentiation

iPSCs were differentiated into mDA neurons as previously described.^35,36^ In short, iPSC were cultured in 1:1 Neurobasal:DMEM/F12 [Thermo Fisher Scientific], supplemented with N2 supplement 100X [1:100, Thermo Fisher Scientific], B27 supplement 50X [1:50, Thermo Fisher Scientific], 1 % L-Glutamine [200 mM, Invitrogen], 1 % P/S, Y27632 (only day 0) [0.5 µM, Cambridge Bioscience], SB431542 [10 µM, Cambridge Bioscience], LDN193187 [100 nM, Sigma], CHIR99021 [0.9 µM, Tocris Bioscience], Recombinant modified human Sonic Hedgehog C24II (SHH) [200 ng/ml, R&D Systems], Purmophamine (from day 2) [0.5 µM, Cambridge Bioscience]). EBs were plated at day 4 onto Poly-L-ornithine (PO) (15 µg/ml, Sigma), Fibronectin (FN) (5 µg/ml, Invitrogen) and Laminin (LN2020) (5 µg/ml, Sigma) coated wells in 1:1 Neurobasal:DMEM/F12 [Thermo Fisher Scientific], N2 supplement 100 X [1:200, Thermo Fisher Scientific], B27 supplement 50X [1:100, Thermo Fisher Scientific], 1 % L-Glutamine [200 mM, Invitrogen], 1 % P/S, and supplemented between day 0 to 9 with SB431542 (withdrawn at day 6) [10 µM, Cambridge Bioscience], LDN193187 [100 nM, Sigma], CHIR99021 [0.9 µM, Tocris Bioscience], Recombinant modified human Sonic Hedgehog C24II (SHH), [200 ng/ml, R&D Systems], Purmophamine, [0.5 µM, Cambridge Bioscience]). On day 11, cells were drop plated on PO/FN/LN2020 coated plates in Neurobasal [Thermo Fisher Scientific], N2 supplement 100 X [1:200, Thermo Fisher Scientific], B27 supplement 50X [1:100, Thermo Fisher Scientific], 1 % L-Glutamine [200 mM, Invitrogen], 1 % P/S, supplemented with Ascorbic Acid (0.2 mM, Sigma) and Brain-Derived Neurotrophic Factor BDNF (20 ng/ml, Miltenyi Biotech). On day 14, Glial cell-Derived Neurotrophic Factor GDNF (20 ng/ml, Miltenyi Biotech) and N6,2’-O-Dibutyryladenosine 3’,5’-cycle monophosphate sodium salt [db-cAMP (0.5 mM, Sigma)] were added. On day 30, cell were replated and cultured until day 65 of differentiation in medium supplemented with DAPT (2.5 µM, Tocris Bioscience).

### Immunocytochemistry and imaging

iPSC and iPSC-derived dopaminergic neurons were washed and fixed in 4 % paraformaldehyde at RT for 10 min. Cells were then blocked in blocking solution (FBS, 10 % FBS, 0.1 % - 0.3% Triton X-100 [Sigma]) at RT for 1 h, followed by primary incubation overnight at 4°C. Primary antibodies used included OCT4 (Santa Cruz), NANOG (Millipore), TRA-1-60 (Santa Cruz), TRA-1-81 (Millipore), FOXA2 (BD Pharmigen^TM^), LMX1A (Millipore), TH (Aves Labs), MAP2 (Sigma), GIRK2 (Alomone labs), NeuN (Millipore), SYP (Sigma), ZIP14 (Zoa lab, Arizona), ZIP8 (Sigma), ZnT10 (Abcam), NCAM1 (Cell Signaling), Calnexin (Santa Cruz), TOMM20 (Merck), LAMP1 (Merck), GPP130 (BioLegend), cCASP3 (Cell Signaling). The following day, cells were washed 3 times in PBS and incubated with secondary antibodies diluted in blocking solution at RT for 45 min. Secondary antibodies (all from Life Technologies) included Alexa Fluor® 594 Goat Anti-chicken IgG, Alexa Fluor® 488 Goat Anti-mouse IgG, Alexa Fluor® 488 Goat Anti-rabbit IgG, Alexa Fluor® 594 Goat Anti-rabbit IgG, Alexa Fluor® 594 Goat Anti-mouse IgG, Alexa Fluor® 633 Goat Anti-mouse IgG, Alexa Fluor® 594 Goat Anti-mouse IgM. DAPI was used for nuclear staining unless otherwise stated. Cells plated on slides were mounted with ProLong Gold Antifade Mountant (Invitrogen). Images were acquired on an Olympus IX71 inverted TC scope or on a LSM710 Zeiss confocal microscope. Image analysis was performed using ImageJ software.

### qRT-PCR analysis

Total RNA was extracted using the RNeasy kit (QIAGEN). RNA was purified using the DNase I kit (Invitrogen) and cDNA was generated using the SuperScript III Reverse transcriptase (Invitrogen). cDNA was mixed in a qRT-PCR plate with MESA BLUE qPCR 2x MasterMix Plus for SYBR assay (Eurogentec) and qRT-PCR performed on a StepOnePlus^TM^ real-Time PCR System (Applied Biosystems). Gene expression was analysed using the ΔΔCT method, with control 1 mDA line as internal control and GAPDH as the housekeeping gene. Gene expression was measured in the following genes: COX-I, COX-II, and SDHA, and GAPDH.

### Immunoblotting

Cells were resuspended and lysed in RIPA lysis and extraction buffer (ThermoFisher Scientific) containing 1x Protease and Phosphatase Inhibitor cocktail (ThermoFisher Scientific) for 30 min at 4°C. Extracts were then centrifuged at 13.000 rpm, 4°C, for 15 min. The supernatant was collected and protein concentration measured using Pierce^TM^ BCA Protein Assay Kit (ThermoFisher Scientific). Protein absorbance was read at 562 nm using a Spectramax i3x Microplate reader (VWR). 10 µg of total proteins were denatured at 95°C for 5 min in 100 mM dithiothreitol (DTT) and 1x Laemmli buffer (Bio-Rad). Proteins mixtures were separated using a Mini-PROTEAN Tetra Vertical Eletrophoresis Cell for Mini Precast Gels apparatus (Bio-Rad) on 4-20 % Mini-PROTEAN TGX^TM^ Stain-Free Protein Gels (Bio-Rad) and 1x Tris/Glycine/SDS buffer (Bio-Rad). Proteins were then transferred onto a Trans- Blot Turbo^TM^ Mini PVDF Transfer membrane (Bio-Rad) using a Trans-Blot Turbo^TM^ Transfer System (Bio-Rad). Membranes were blocked in 5 % milk in Tris Buffered Saline (TBS, Sigma) with 0.1 % Tween 20 (Sigma) for 1 h at RT. Membranes were incubated with primary antibodies in 1 % milk in TBS – 0.1 % Tween 20 overnight at 4°C with constant gentle shaking. Primary antibodies included cCASP3 (Cell Signaling), GPP130 (BioLegend), JNK (Cell Signaling), P-PNK (Cell Signaling) and β-actin (Sigma). Membranes were then incubated with appropriate horseradish peroxidase-conjugated antibody secondary antibody (HRP-conjugated Anti-Rabbit IgG and Anti-Mouse IgG, Cell Signaling) in 1 % milk in TBS – 0.1 % Tween 20 for 1 h at RT. Membranes were visualized with ChemiDocTM MP (Bio-Rad), using Clarity Western ECL Substrate (Bio-Rad). ImageJ software (NIH) was used for protein quantification, and normalization performed against β-actin.

### ICP-MS analysis

Cell pellets from day 65 neuronal cultures were digested in 1 ml of 3 % Nitric acid (Millipore) on a shaker at 85°C overnight followed by an incubation of 2 h at 95°C the following day. Samples were centrifuged for 10 min at 15.000 x g to remove any remaining cellular debris. The metal ion isotopes ^55^Mn, ^56^Fe, ^66^Zn, and ^44^Ca, were measured by triplicate using an Agilent 7500ce ICP-MS instrument with collision cell (in He mode) and Integrated Autosampler (I- AS) using ^72^Ge as internal standard. The following experimental parameters were used: a) plasma: RF power 1500 W, sampling depth 8.5 mm, carrier gas 0.8 L/min, make-up gas 0.11 L/min; b) quadrupole: mass range 1-250 amu, dwell time 100 msec and 0.1 sec/point integration time. Protein concentration for each sample was previously determined using the BCA Protein Assay Kit (ThermoFisher Scientific) to be used for normalization purposes.

### Pulse-chase assay

iPSC-derived neurons at day 65 were treated with 100 µM MnCl_2_ (Sigma) for 48 h (pulse phase). Neurons were then washed 5 times in HBSS without Ca and Mg (Invitrogen) and incubated with 1 ml HBSS without Ca and Mg for 1 h (chase phase). Supernatant and cell pellets were collected separately. Cell pellets were digested in 3 % Nitric acid as described above. The metal ion isotopes Mn-55, Fe-56, Zn-66, Ca-44, and Cd-111 were measured by an Agilent 7,000 Series ICP-MS in Helium collision mode in both digested samples and supernatants.

### Mitochondrial membrane potential measurement

Mitochondria membrane potential was measured using the red-fluorescent probe tetramethylrhodamine-methyl ester (TMRM) following manufacturer’s recommendations (Abcam). In short, mDA neurons at day 30 were plated onto PO/FN/Lam coated 35 mm round FluoroDish^TM^ (World Precision Instrument) and grown until reaching maturity (day 65). On the day of experiment, cells were washed in PBS then incubated for 30 min at 37°C in Neurobasal phenol-free working solution supplemented with TMRM 20 nM (Abcam), Calcein 4 µM (Insight Biotechnology), and Hoechst 2 µM (ThermoFisher). Staining medium was then removed and cells bathed in Neurobasal phenol-free supplemented with TMRM 20 nM for live cell imaging. Images were acquired on a CSU-22 Spinning Disk Confocal (Zeiss), using the Volocity 6.0 software. Image processing and quantification of staining intensity was performed on Fiji using a home-made macro (Dr. Dale Moulding, UCL GOS ICH, London). This macro measures staining intensity in mitochondria by identifying mitochondria area size using the “tubeness” mode, measuring total volume and total intensity of pixels within identified volume. Total intensity was divided by total volume to provide average intensity across each image.

### Mitochondrial respiration

Mitochondrial functions were assessed by directly measuring the oxygen consumption rate (OCR) of mature neurons on a Seahorse XFe and XF extracellular Flux Analyzer (Agilent Technologies) as per manufacturer’s recommendations. In short, mDA neurons at day 30 of differentiation were seeded at high density (>50,000 cells/well) in a PO/FN/Lam coated 96 wells Seahorse XF cell culture microplate (Agilent Technologies) and cultured for a further 35 days until full neuronal maturation. On the day prior to assay, the sensor cartridge was hydrated (XFe 96 FluxPak, Agilent Technologies) with the Seahorse XF Calibrant (Agilent Technologies) at 37°C in a non-CO_2_ incubator, for a minimum of 24 h. On the day of assay, assay medium was prepared by supplementing the Seahorse XP DMEM medium, pH 7.4 (Agilent Technologies) with 1 mM pyruvate (Sigma), 2 mM glutamine (Sigma), and 10mM glucose (Sigma) and warmed to 37°C in a non-CO_2_ incubator for 45 min to 1 h prior to assay. Oligomycin (final concentration: 2 µM, Sigma), FCCP (final concentration: 1 µM, Sigma), Antimycin-A (final concentration: 2.5 µM, Sigma) prepared in Seahorse XP DMEM assay medium were sequentially injected (port A – oligomycin, port B – FCCP, port C – FCCP, Port D – Antimycin-A) into the microplate to modulate distinct components of the ETC and to directly measure parameters of mitochondrial bioenergetics. After OCR data was obtained, in plate protein extraction using RIPA buffer and BCA protein quantification were performed. Data analysis was performed on the Wave 2.6 software (Agilent Technologies) and OCR was normalised to protein concentration in each well.

### Uptake assay using FM^TM^ 1-43

The FM™ 1-43 (N-(3-Triethylammoniumpropyl)-4-(4-(Dibutylamino) Styryl) Pyridinium Dibromide) (Invitrogen) membrane probe was used to identify actively firing neurons and to investigate mechanisms of activity-dependent vesicle cycling. Prior to assay, stock solutions of Potassium Chloride (KCl, 1.375 M, Sigma), Magnesium Chloride (MgCl_2_, 1.5 M, Sigma), and Calcium Chloride (CaCl_2_, 0.74 M, Sigma) were prepared. Working solutions were prepared on day of assay. Solution 1: HBSS with Calcium and Magnesium supplemented with MgCl_2_ (2 mM) and CaCl_2_ (2 mM). Solution 2: HBSS with Calcium and Magnesium supplemented with MgCl_2_, CaCl_2_ (2 mM), KCl (60 mM), NBQX 10 µM (Sigma), FM^TM^ 1-43 dye (5 µg/ml). Solution 3: HBSS with Calcium and Magnesium supplemented with MgCl_2_ (2 mM), CaCl_2_ (2 mM) and FM^TM^ 1-43 dye (5 µg/ml). Cells grown on Nunc Lab-TekTM II chamber slides (ThermoFischer) were washed with HBSS and incubated with solution 1 for 10 min at 37°C. Medium was then replaced with solution 2 and cells incubated for 2 min at 37°C. Medium was replaced with solution 3 and cells incubated for 15 min at 37°C. Cells were washed 3 times with HBSS without Calcium and Magnesium and fixed 10 min at RT with 4 % PFA in HBSS without Calcium and Magnesium. Slides were mounted with ProLong Gold Antifade Mountant (Invitrogen). Images were acquired on the LSM710 Zeiss confocal microscope. Images were analysed using a home-made macro in Fiji (Dr. Dale Moulding, UCL GOS ICH, London). This macro measures the total surface area of imaged neurites as well as measures the intensity of staining by FM™ 1-43 dye. The macro then measures the ratio between total surface and staining intensity to provide an average fluorescence intensity unit per image.

### Calcium imaging using Calbryte^TM^ 520

Neurons at day 30 of differentiation were plated on PO/FN/Lam coated 35 mm round FluoroDish^TM^ (World Precision Instrument) and grown until reaching maturity (day 65). On the day of experiment, cells were washed twice in Krebs solution (NaCl 7.03 g/L, KCl 0.44 g/L, NaHCO_3_ 1.3 g/L, Glucose 2.07 g/L, MgCl_2_ 0.245 g/L, NaH_2_PO_4_ 0.187 g/L and CaCl_2_ 0.3675 g/L in deionised water), then incubated at 37°C in a CO_2_ incubator for 20 min with Calbryte^TM^ 520 AM (AAT Bioquest) at final concentration of 1 µM and supplemented with 0.02 % of Kolliphore® EL (Sigma) in Krebs solution. Following incubation, cells were washed three times in Krebs solution for 10 min at 37°C. Ca^2+^ transients were also measured following addition of Tetrodotoxin (TTX) (10 µM, Sigma) with NBQX (2,3-dioxo-6-nitro-7-sulfamoyl-benzo[f]quinoxaline) (10 µM, Sigma) for 120 s. The purpose was to block Na+ channels and AMPA receptor (α-amino-3-hydroxy-5- methyl-4-isoxazolepropionic acid receptor), respectively. Fluorescence imaging was performed on an Olympus BX51 microscope equipped with a 40x water dipping lens (LUMPLFLN40xW, NA 0.8, Olympus Europa), and an EMCCD camera (iXon Ultra 897, Andor Technology). Calbryte 520 was excited at 470nm using an OptoLED (Cairn Research Limited), and fluorescence emission was collected at 525/50nm. Images (512X512 pixels2) were acquired sequentially at 2Hz for a period of 2-minutes. Data analysis was performed using the ImageJ software.

### Bulk-RNA sequencing

Bulk-RNA sequencing was performed as previously described.^36^ In short, total RNA of 66 samples (2 controls, 6 patient lines and 3 isogenic controls in 3 biological replicates, unexposed or exposed with MnCl_2_ for 48 h) were extracted using the RNeasy mini kit (Qiagen) following manufacturer’s recommendations. RNA libraries were prepared from 100 ng of total RNA using the Kapa mRNA Hyper Prep kit (Roche) and sequenced with a Novaseq SP v1.5 (100 cycles, 44 M reads/samples single reads) as per manufacturers recommendations and performed at UCL Genomics, GOS ICH Zayed Centre for Research. FASTQ files were loaded onto Galaxy web platform (www.usegalaxy.org) and public server of Galaxy used for further analysis. Quality control (QC) was performed using the FastQC v0.11.9 and MultiQC v1.11 modes in Galaxy. Reads were mapped to a human reference genome (GRCH38) using HISAT2 v2.2.1. Count genes were mapped to respective gene using FeatureCounts v.2.0.1 excluding multiple mapping, duplicates, and chimeric fragments. Differentially expressed genes were determined using EdgeR v.3.36.0, with a threshold set for P value 1 for statistical significance, and filtering low counts at 0.35 minimum counts per million. Comparisons were performed between controls and patients unexposed, controls unexposed and controls manganese-exposed, controls manganese-exposed and patients manganese-exposed, controls unexposed and patients manganese-exposed. Gene ontology enrichment was performed using ShinyGO 0.76.2 for the biological processes and ClueGo v2.5.8 + CluePedia v1.5.8 for cellular components, molecular processes and reactome networks, with Benjamini-Hochberg P-value correction of false discovery rate (FDR)<0.05. Results from the expression analysis along with the raw sequence data were deposited in GEO (Gene Expression Omnibus).

### Statistical analysis

Statistical analysis was performed using the GraphPad Prism v9.4.1 software. Samples were compared using the Student’s unpaired two tailed t-test for simple comparisons.^36,38^ Results are reported as mean ± standard error or the mean (SEM) from at least three independent biological replicates. Significant levels were determined by P-value and are represented by asterisks. P-values are represented as (*)P=0.05-0.01, (**)P=0.01-0.001, (***)P<0.001.

### Data availability

The authors confirm that the raw transcriptomics data supporting the findings of this study are available within the article and its supplementary material. These data will be made openly available in Gene Expression Omnibus (GEO) upon request.

---

**Figure S1.**
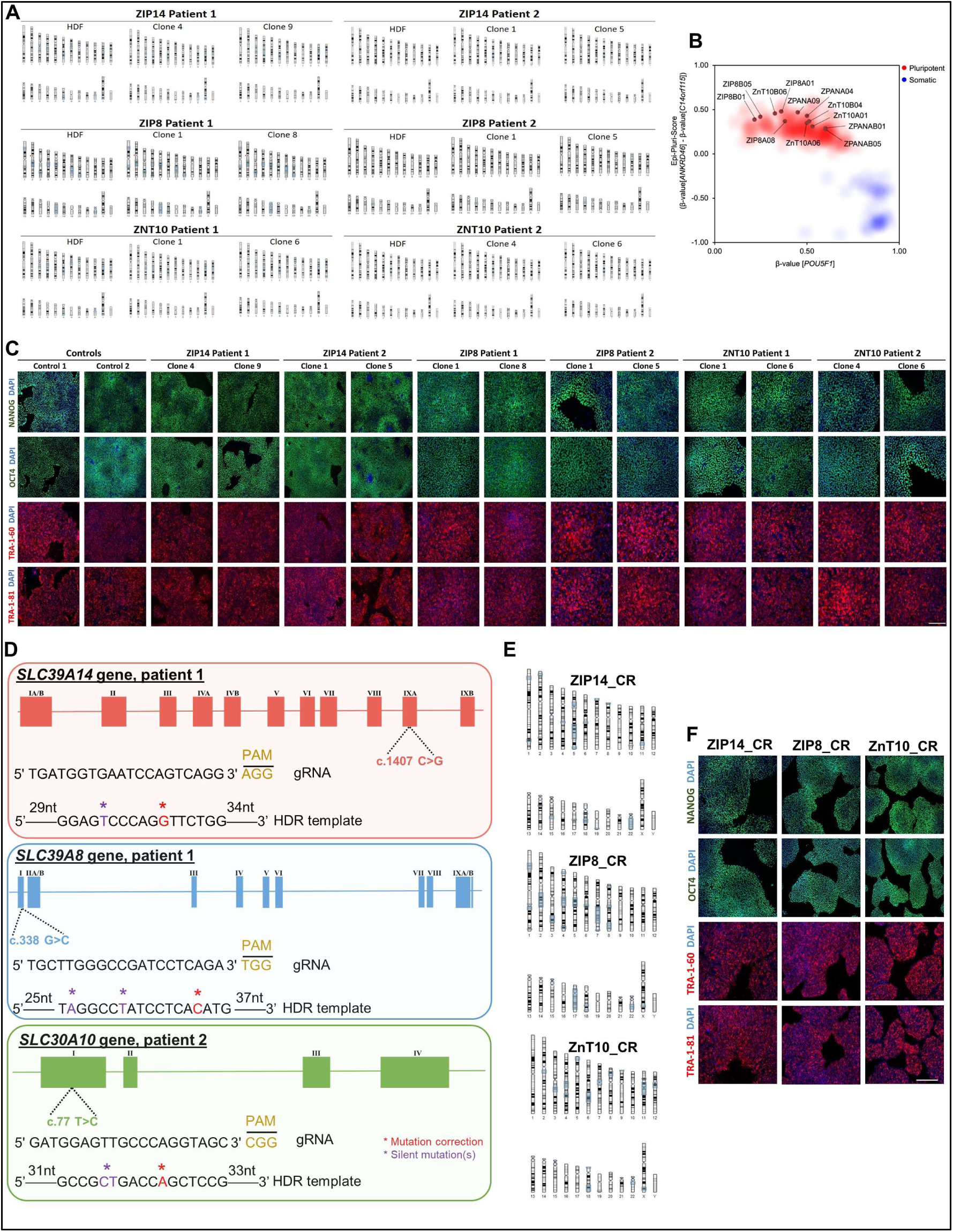
iPSC characterization and generation of CRISPR-Cas9 isogenic controls related to Figure 1. **(A)** SNP analyses of the iPSC lines generated in this study show normal karyotype, without significant differences to derived fibroblast line. Two clones per line were initially characterized and compared to the original human dermal fibroblast (HDF). **(B)** Epi-Pluri-Score analysis of iPSC lines show all generated lines cluster within the pluripotent stem cell lines (red), rather than with somatic cell lines (blue). Analysis generated by Cygenia, Epigenetic Diagnostics, Aachen, German. **(C)** Immunofluorescence analysis for pluripotency markers OCT4, NANOG, TRA-1-60, and TRA-1-81 show all iPSC lines express these pluripotent markers. Scale bar, 100 µm. **(D)** gRNA and HDR repair template design for CRISPR-Cas9 single nucleotide correction in the *SLC39A14* patient 1, *SLC39A8* patient 1, and *SLC30A10* patient 2 iPSC lines. **(E)** SNP analyses of the CRISPR-corrected iPSC lines (ZIP14 CR02, ZIP8 CR04, and ZnT10 CR04) show normal karyotype, without significant differences to derived fibroblast line. **(F)** Immunofluorescence analysis for pluripotency markers OCT4, NANOG, TRA-1-60, and TRA-1-81 show all isogenic control lines express these pluripotent markers. Scale bar, 100 µm.

**Figure S2.**
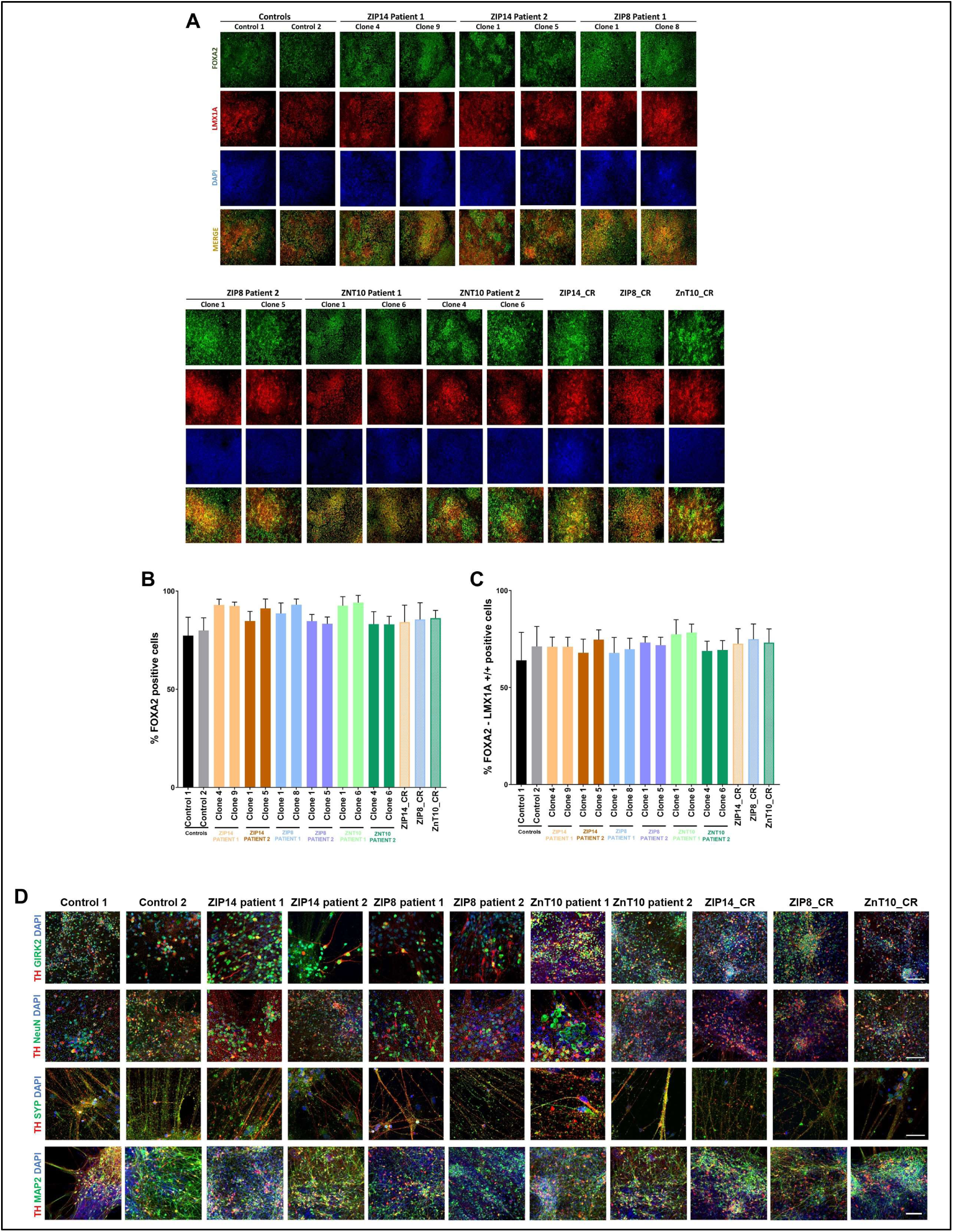
Characterization of iPSC-derived mDA neuronal identity at precursor (day 11) and mature stage (day 65), related to Figure 1. **(A)** Images of day 11 neuron precursors stained for midbrain progenitor markers FOXA2 and LMX1A. **(B-C)** Quantitative analysis of DAPI/FOXA2 positive cells at day 11 of mDA differentiation (B) and quantitative analysis of FOXA2/LMX1A positive cells at day 11 of mDA differentiation (C). Comparable to the literature, there is a high percentage of cells double-positive for FOXA2/LMX1A, suggesting an acceptable level of mDA neuronal progenitors. Values are given as means ± SEM. **(D)** Immunofluorescence analyses for neuronal maturity markers GIRK2, NeuN, SYP, and MAP2 together with mDA marker TH. Here are shown selected clones for each patient lines that were carried forward for downstream experiments, ZIP14 patient 1 clone 9, ZIP14 patient 2 clone 1, ZIP8 patient 1 clone 1, ZIP8 patient 2 clone 1, ZnT10 patient 1 clone 1, and ZnT10 patient 2 clone 6 were selected, Scale bar, 100 µm.

**Figure S3.**
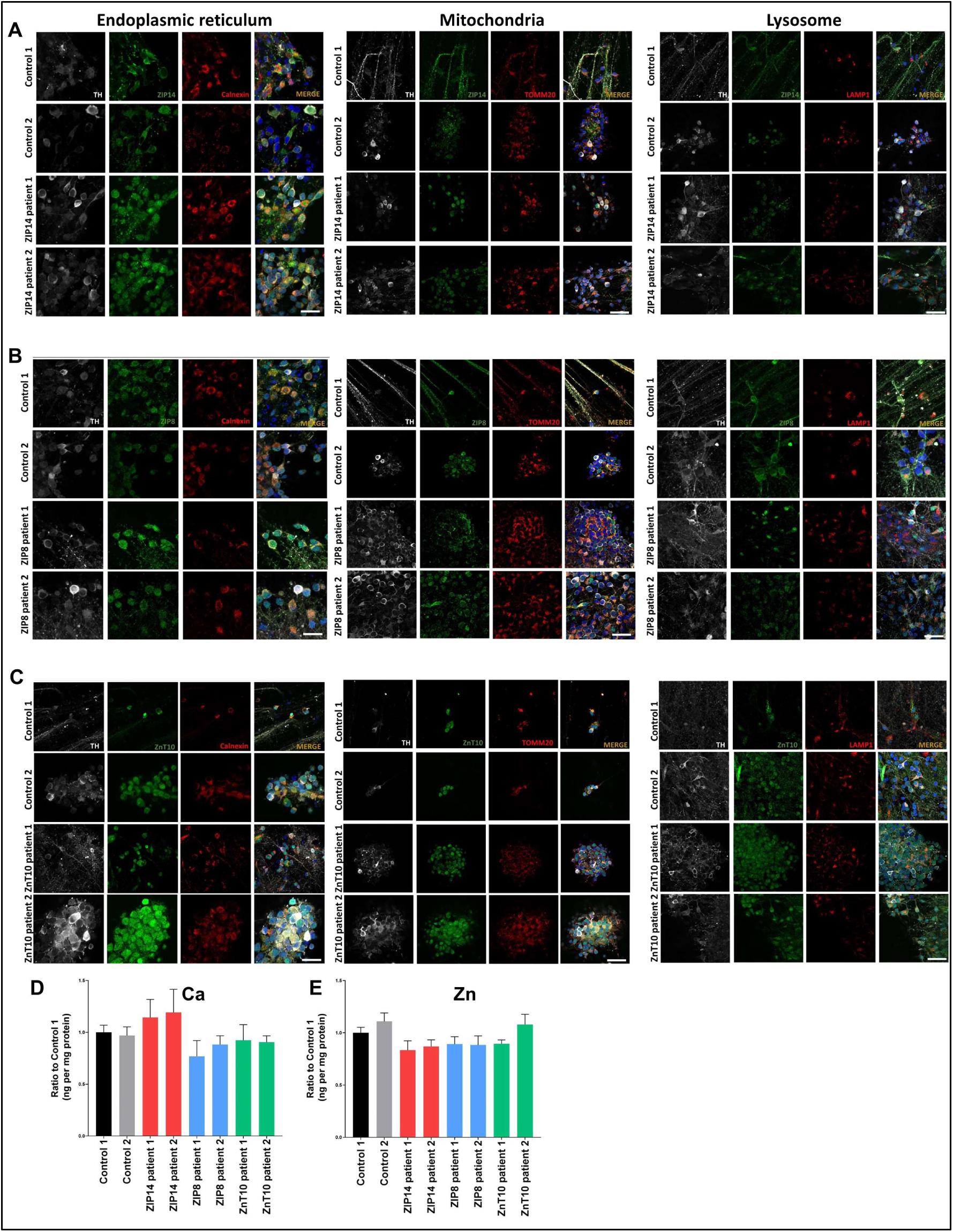
Subcellular localization of metal-ion transporters ZIP14, ZIP8, and ZnT10 in mDA neurons and Ca and Zn intracellular levels, related to Figure 1. **(A-C)** Immunofluorescence for the endoplasmic reticulum (calnexin), mitochondria (TOMM20), and lysosomes (LAMP1) in TH-positive neurons show co-localization of ZIP14, ZIP8, and ZnT10 with the ER, mitochondria, and nucleus (DAPI), but not with lysosomes. Scale bars, 20 µm (ER) and 50 µm (TOMM20, LAMP1). **(D-E)** ICP-MS analysis for intracellular levels of calcium (D) and zinc (E) do not show significant intracellular differences. Values are given as means ± SEM.

**Figure S4.**
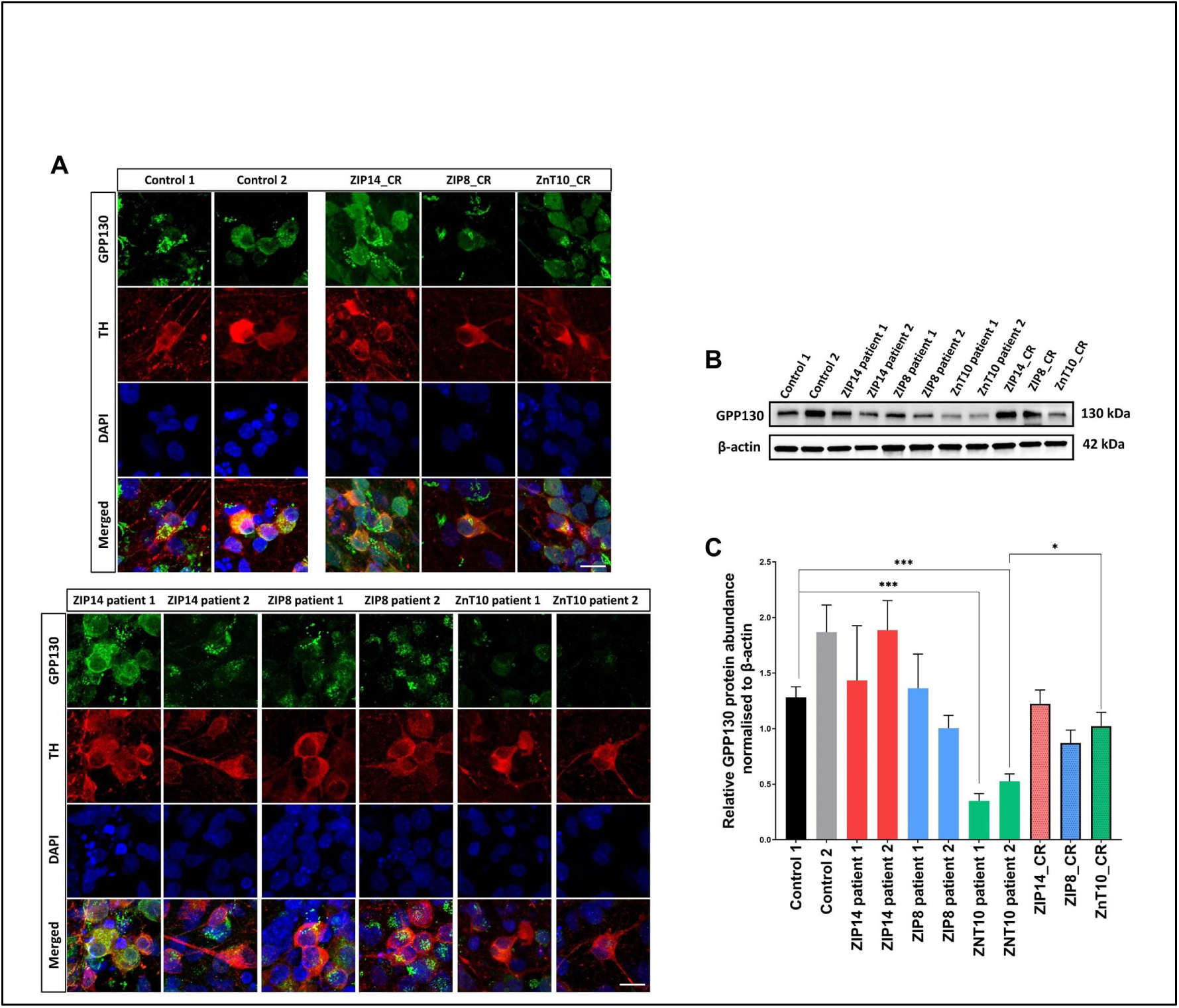
Immunofluorescence for the manganese-specific sensor GPP130 for the efflux activity of ZnT10 transporter, related to Figure 1. **(A)** Day 65 neurons were stained for the Golgi marker GPP130 and mDA marker TH, which show a reduction in GPP130 signal intensity in the ZnT10 patient lines. Scale bar, 10 µm. **(B-C)** Immunoblot of total GPP130 protein (130 kDa) with β-actin (45 kDa) as housekeeping gene (B). Relative abundance of GPP130 protein, normalized to β-actin. (^∗^p=0.05-0.01, **p=0.01-0.001, p***< 0.001, unpaired Student’s t test). N= 4 – 10 independent experiments. Values are given as means ± SEM.

**Figure S5.**
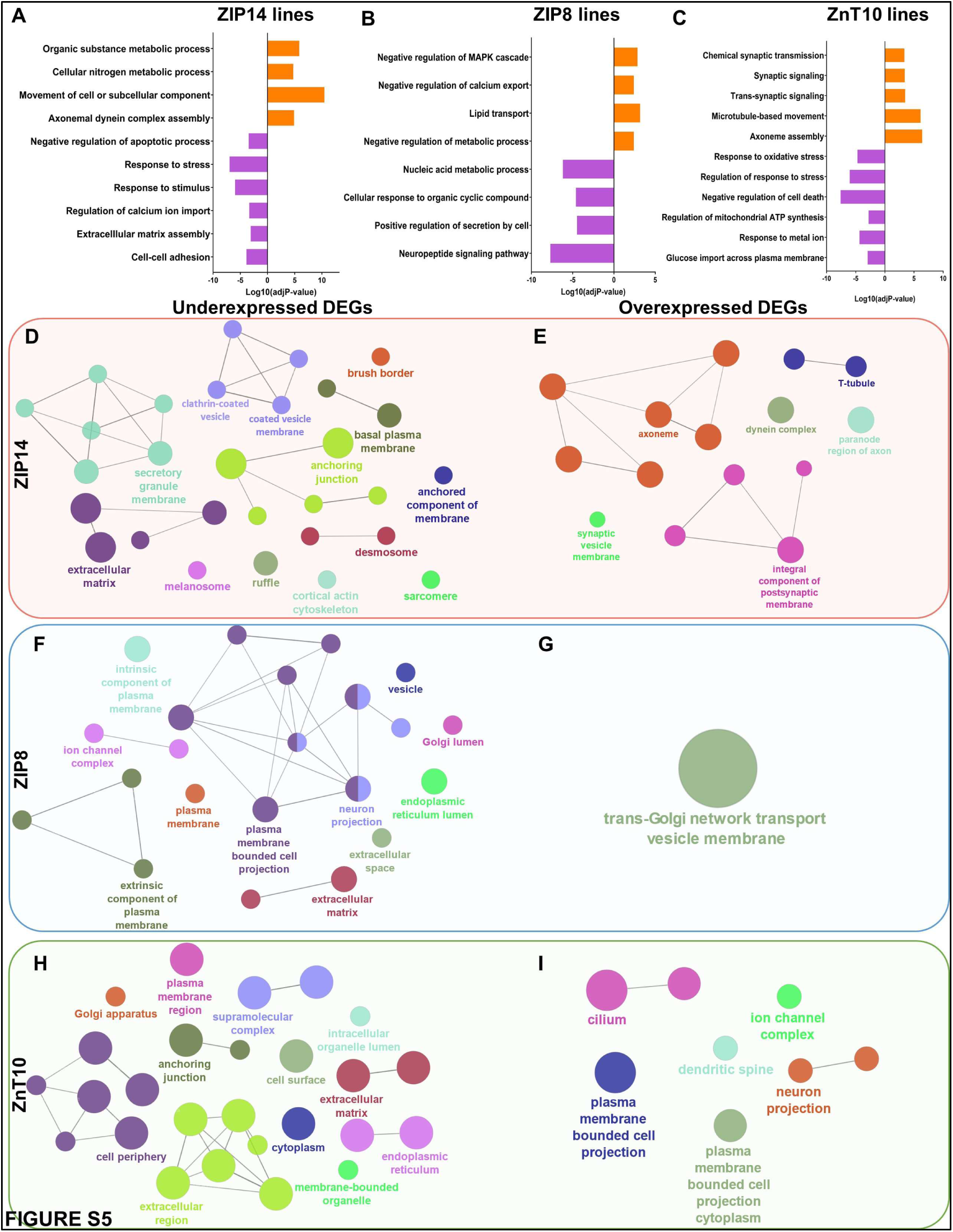
Dysregulation of biological processes and cellular components in ZIP14, ZIP8, and ZnT10 lines, related to Figure 2 and Supplementary Table 1. **(A-C)** Gene ontology (GO) terms enrichment for biological process of underexpressed (blue) and overexpressed (red) protein coding genes in ZIP14 (A), ZIP8 (B), and ZnT10 (C) lines. **(D-I)** ClueGO analysis of GO terms enrichment for underexpressed and overexpressed DEGs, showing nodes network of cellular components in ZIP14 (D-E), ZIP8 (F-G), and ZnT10 (H-I) lines. Network graph nodes represent GO terms and edges (connections) indicate shared genes between GO terms. Colour coding represents functional groups of GO terms. Only the GO functional groups exhibiting higher statistically significant differences, using Benjamini-Hochberg P-value correction (FDR<0.05) are shown.

**Figure S6.**
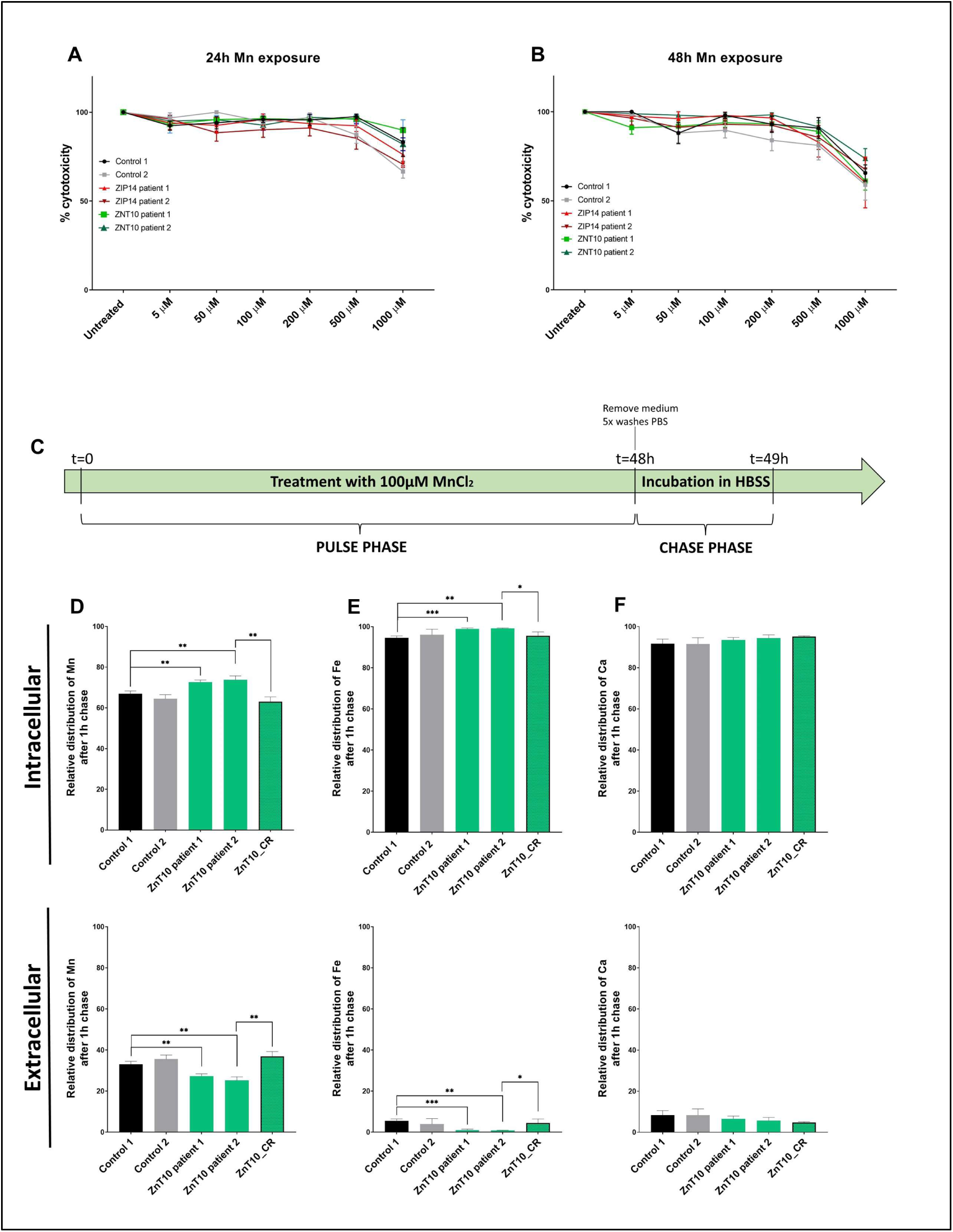
Pulse chase assay confirms manganese efflux activity of ZnT10 related to Figure 3. **(A-B)** MTT assay confirms that exposure to 100 um MnCl_2_ for 48h does not cause significant cell death in mDA neurons. **(C)** Schematic of pulse-chase assay: neurons are treated for 48 h with 100uM MnCl_2_ (pulse phase), then incubated for 1 h in HBSS without manganese. **(D-F))** Relative distribution of manganese (B), iron (C), calcium (D), and cadmium (E) between intracellular and extracellular compartments. (^∗^p=0.05-0.01, **p=0.01-0.001, p***< 0.001, unpaired Student’s t test). Values are given as means ± SEM.

**Figure S7.**
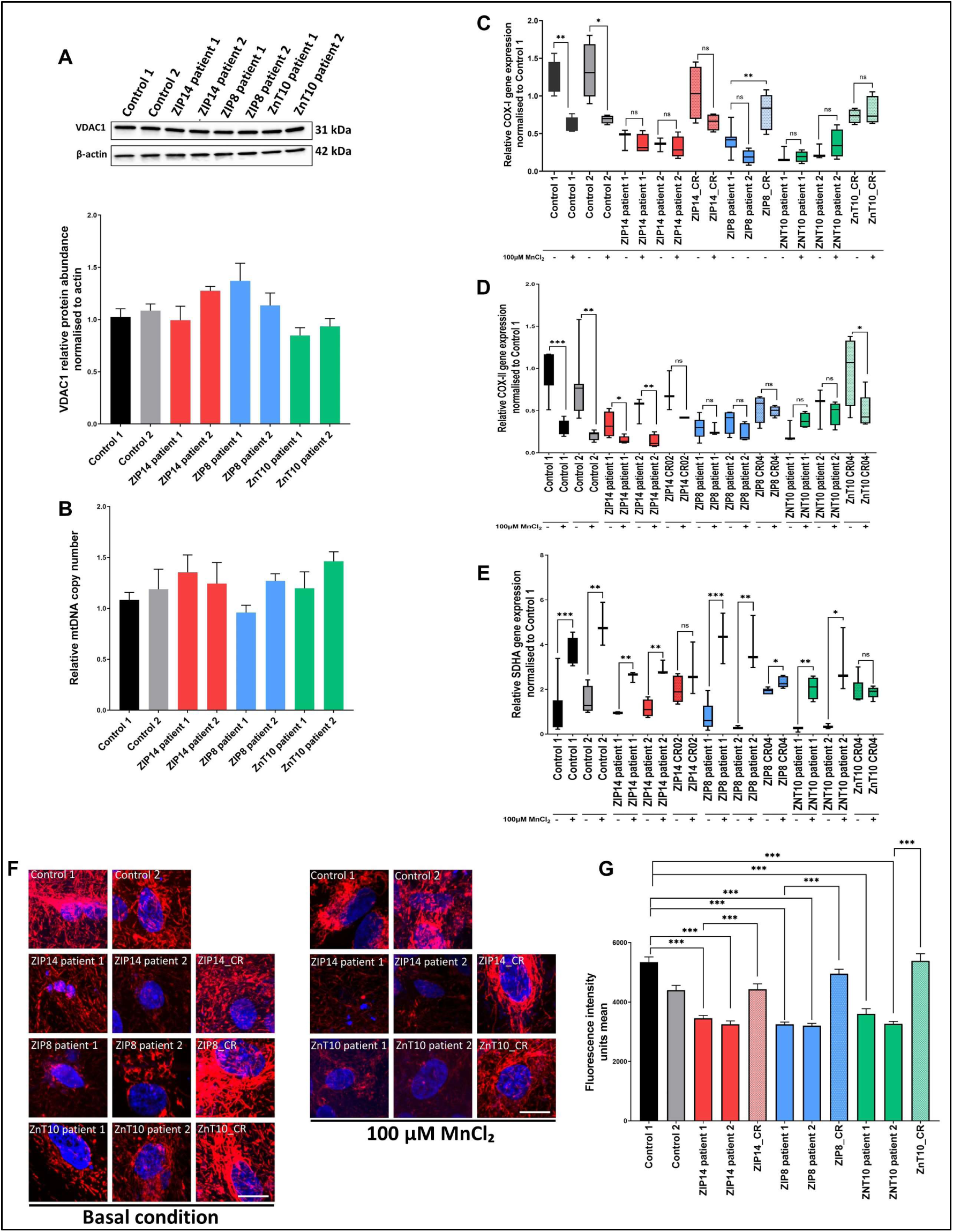
Mitochondrial integrity and gene expression , related to Figure 5. **(A)** Immunoblot analysis for VDAC1 (A) and relative quantification (B) shows no difference in mitochondrial mass between lines. N=3 – 7 independent experiments. Values are given as means ± SEM. **(B)** qRT-PCR analysis for mtDNA copy number. N=3 independent experiments. Values are given as means ± SEM. **(C-E)** qRT-PCR analyses for COX-I (C), COX-II (D), and SDHA (E), relative to GAPDH and normalized to an internal control. (^∗^p=0.05-0.01, **p=0.01-0.001, p***< 0.001, unpaired Student’s t test). Box-and-whisker plot shows median with min to max values. Values are given as means ± SEM. **(F)** Representative zoomed-in TMRM immunostaining in mature neurons, in both physiological and manganese exposed conditions. Scale bar = 30 µm. **(G)**TMRM fluorescence intensity measurements in basal and Mn-exposed conditions, between control and patient lines. (^∗^p=0.05-0.01, **p=0.01-0.001, p***< 0.001, unpaired Student’s t test). Box-and-whisker plot shows median with min to max values. Values are given as means ± SEM.

**Figure S8.**
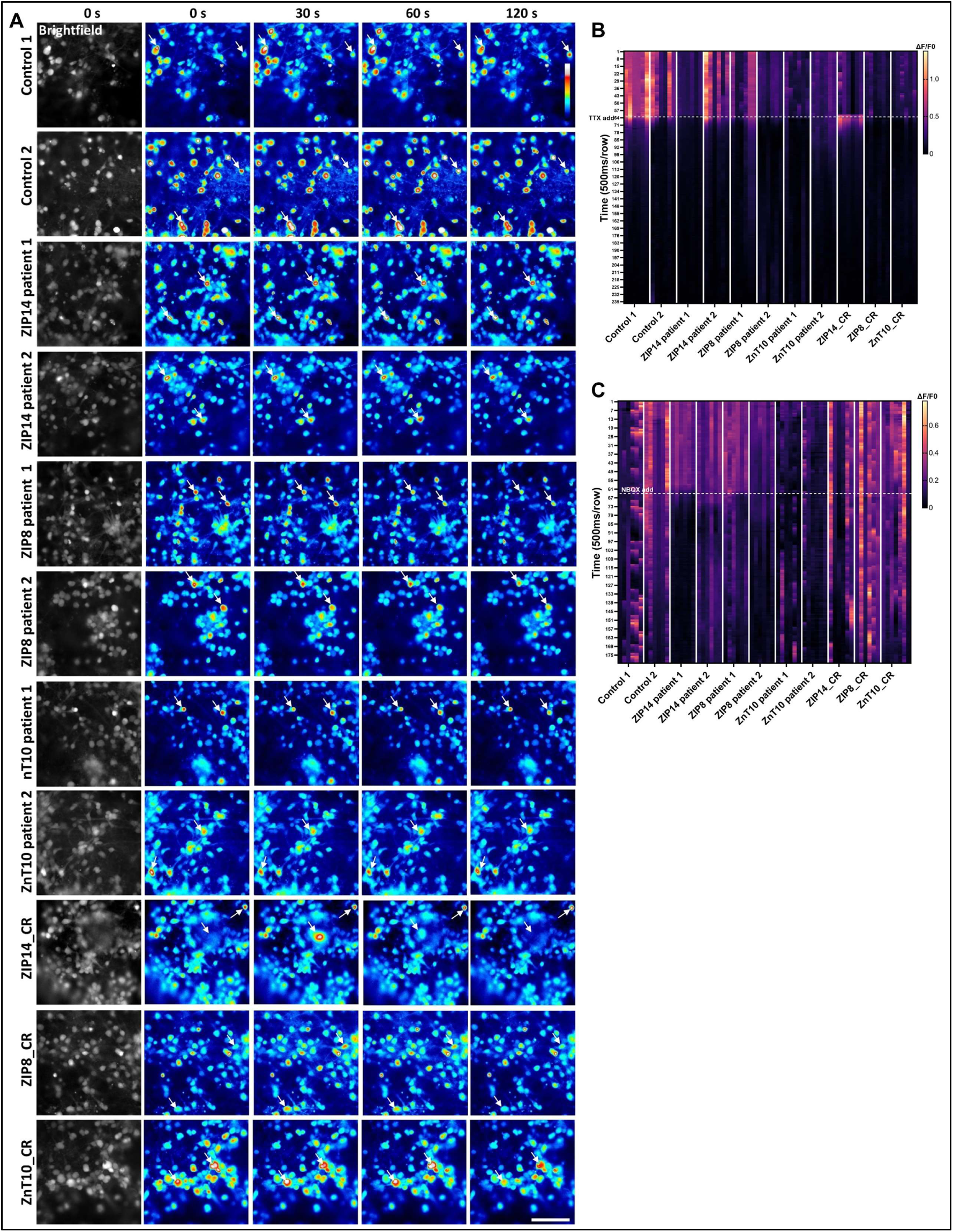
Measurement of calcium signalling in control and patient lines, and following addition of modulators of neuronal excitability, related to Figure 7. **(A)** Representative Ca^2+^ transients over a 2 min period in neurons labelled with Calbryte 520 AM. Arrows indicate cells with change in Ca^2+^ fluxes over time (red to blue: high to low fluorescence intensity), **(B)** Ca^2+^ transients measured following addition of 10 µM TTX, for a total of 120 s. Scale represents ΔF/F0 fluorescence intensity, ranging from 0 (low ΔF/F0, dark purple) to 0.8 (high ΔF/F0, yellow). n=6 representative cells in one experiment. **(C)** Ca^2+^ transients measured following addition of 10 µM NBQX, for a total of 120 s. Scale represents ΔF/F0 fluorescence intensity, ranging from 0 (low ΔF/F0, dark purple) to 0.8 (high ΔF/F0, yellow). n=6 representative cells in one experiment

